# HSD2 neurons are evolutionarily conserved and required for aldosterone-induced salt appetite

**DOI:** 10.1101/2024.04.24.590990

**Authors:** Silvia Gasparini, Lila Peltekian, Miriam C. McDonough, Chidera J.A. Mitchell, Marco Hefti, Jon M. Resch, Joel C. Geerling

**Affiliations:** Department of Neurology, University of Iowa; Department of Neuroscience and Pharmacology, University of Iowa; Department of Pathology, University of Iowa; Iowa Neuroscience Institute, University of Iowa

## Abstract

Excessive aldosterone production increases the risk of heart disease, stroke, dementia, and death. Aldosterone increases both sodium retention and sodium consumption, and increased sodium consumption predicts end-organ damage in patients with aldosteronism. Preventing this increase may improve outcomes, but the behavioral mechanisms of aldosterone-induced sodium appetite remain unclear. In rodents, we identified aldosterone-sensitive neurons, which express the mineralocorticoid receptor and its pre-receptor regulator, 11-beta-hydroxysteroid dehydrogenase 2 (HSD2). Here, we identify HSD2 neurons in the human brain and use a mouse model to evaluate their role in aldosterone-induced salt intake. First, we confirm that dietary sodium deprivation increases aldosterone production, HSD2 neuron activity, and salt intake. Next, we show that activating HSD2 neurons causes a large and specific increase in salt intake. Finally, we use dose-response studies and genetically targeted ablation of HSD2 neurons to show that aldosterone-induced salt intake requires these neurons. Identifying HSD2 neurons in the human brain and their necessity for aldosterone-induced salt intake in mice improves our understanding of appetitive circuits and highlights this small cell population as a therapeutic target for moderating dietary sodium.

## Introduction

Aldosterone regulates cardiovascular function and electrolyte homeostasis. Its production rises during hypovolemic states, including prolonged sodium deprivation. A primary function of this adrenal steroid hormone is to maintain blood volume by increasing sodium reabsorption (1, 2).

Up to 20% of patients with hypertension have primary aldosteronism, a condition characterized by excessive production of aldosterone (3). Relative to others with hypertension, patients with aldosteronism suffer from stroke, myocardial infarction, atrial fibrillation, and heart failure at several-fold higher rates (4–6), as well as moderately elevated rates of diabetes and dementia (7–9).

The reasons for these risk discrepancies are not fully understood, and one aspect that has received little attention is salt consumption. In addition to its primary, sodium-retaining effect, aldosterone increases sodium appetite in rats (10–12), and among human patients with hypertension, the threshold for sodium taste perception is higher in those with aldosteronism (13). Despite their excessive sodium retention, patients with aldosteronism occasionally report salt cravings (14–16), and those with aldosterone-secreting adenomas consume the most sodium (17). Excessive salt intake is particularly problematic in aldosteronism because sodium intake predicts the severity of cardiac complications, including left ventricular hypertrophy (18–21). Decreasing salt intake reduces blood pressure and may also reduce cardiovascular complications (19–23), but long-term behavioral recommendations to reduce dietary sodium are notoriously ineffective. Therefore, it may help to determine, and then disrupt, the mechanism of aldosterone-induced salt appetite.

It is now well-established that aldosterone selectively acts in cells that express the HSD2 enzyme (11-beta-hydroxysteroid dehydrogenase type 2) and the mineralocorticoid receptor. HSD2 is a pre-receptor filter that confers selectivity for aldosterone by metabolizing cortisol and other glucocorticosteroids, which otherwise occupy most mineralocorticoid receptors (24, 25). HSD2 makes a mineralocorticoid receptor-expressing cell aldosterone-sensitive because it frees more receptors to bind aldosterone, which circulates at concentrations 100-fold lower than cortisol (26–28). HSD2 expression, in addition to the mineralocorticoid receptor, is thus the defining feature of aldosterone-sensitive cells.

HSD2 expression is limited to one, small population of mineralocorticoid receptor-expressing cells in the rodent brain. These cells, known as the HSD2 neurons, are located in the nucleus of the solitary tract (NTS; 29, 30), which is a hub for interoceptive information arriving from the heart, lungs, gut, and other viscera. Hypovolemic states like dietary sodium deprivation increase aldosterone production and activate the HSD2 neurons (27, 31–34). HSD2 neurons from aldosterone-infused mice fire pacemaker-like action potentials (35), and chemogenetic stimulation of HSD2 neurons increases salt consumption (35, 36). This circumstantial evidence suggests that these cells play a role in aldosterone-induced sodium appetite, but it remains unclear if physiologically relevant levels of aldosterone alter sodium appetite, whether aldosterone or other mineralocorticosteroids induce sodium appetite in mice (37, 38), and if so, whether HSD2 neurons are involved. It also remains unclear if other mammals, particularly humans, have HSD2 neurons.

Previous work on mineralocorticoid-induced salt intake viewed the hindbrain as having an inhibitory role, and previous attempts to identify a site of action focused on forebrain regions that included the lamina terminalis, hypothalamus, and amygdala (39–47). Within the lamina terminalis, some investigators concluded that the subfornical organ controls both thirst and sodium appetite (48, 49) and suggested that aldosterone triggers an intracellular signaling cascade that shifts behavior from thirst to sodium appetite (50). Still others proposed that aldosterone promotes sodium appetite via mineralocorticoid receptors or membrane receptors in the amygdala (39, 41, 46). Importantly, however, none of these forebrain regions express HSD2, and no studies have evaluated the necessity of any hindbrain region for aldosterone-induced salt intake.

In this study, we tested whether the human brainstem contains HSD2 neurons, then used genetically targeted, cell-type-specific methods in mice to test the hypothesis that HSD2 neurons are necessary for aldosterone-induced sodium appetite. After confirming that dietary sodium deprivation activates HSD2 neurons and that activating HSD2 neurons specifically increases sodium appetite, we identified a dose-response relationship between aldosterone and sodium appetite, then used this information to selectively ablate HSD2 neurons or neighboring NTS neurons to evaluate their necessity for aldosterone-induced sodium appetite.

## Materials and Methods

### Human, porcine, and rat brainstem tissue

Human tissue procurement protocols were reviewed by the University of Iowa’s Institutional Review Board (IRB) and determined not to represent human subjects research under the Revised Common Rule. Consent for research tissue donation was obtained from the next of kin by the University of Iowa Department of Pathology in accordance with federal and Iowa law. We obtained the caudal brainstem from 11 human brains removed after death from a variety of non-neurologic causes. In accordance with Iowa law, no tissue from elective terminations was used for this project. The mean (± SD) post-mortem interval between death and autopsy was 33.5 ± 18 hours (range: 15 to 50 hours). Average age at death was 35 ± 29.6 years (range: 33 gestational weeks to 77 years). Sex was female in two cases and male in nine. After fixing each brainstem for 2–3 days at 4 °C in 10% formalin-PBS (SF100-20, Fischer Scientific), we cryoprotected the brainstem in 30% sucrose-PBS for an additional 1–2 days until it sank. Next, we used a freezing-stage microtome to cut 40 µm-thick tissue sections in the transaxial plane. We collected tissue sections into 12 separate, 1-in-12 series from each brainstem, resulting in an approximately 0.5 mm interval between successive tissue sections within each series. We stored each tissue series in cryoprotectant at −30 °C until removal for further processing. We also used this protocol to process caudal brainstem tissue that had been gifted to us from four pigs that were euthanized and autopsied after other research studies at our institution. For rat brainstem tissue, we obtained new, unpublished images from slides prepared for and described in a previous publication (51).

### Mice

All mice were group-housed in a temperature- and humidity-controlled room with a 12/12-hour light/dark cycle. Every mouse had *ad libitum* access to water and standard rodent chow (Envigo 7013) prior to the experiments described below. Some mice were provided low-sodium chow (TD-130591, Teklad/Envigo) for one or more days in experimental protocols described below. In addition to C57BL6/J mice, where specified below, we used hemizygous *Hsd11b2*-Cre or *Th*-IRES-Cre mice maintained on a C57BL6/J background (**Table 1**). All experiments were conducted in accordance with the guidelines of the Institutional Animal Care and Use Committees at the University of Iowa (protocol numbers 00720011 and 3072011).

**Table 1.** Cre-driver mice.

| Strain | References | Source Information | Key Gene |
| --- | --- | --- | --- |
| *Hsd11b2-Cre* | (36) | Jax 030545<br>http://www.informatics.jax.org/allele/key/871609 | IRES-Cre-GFP inserted after the termination codon of *Hsd11b2* |
| *Th-IRES-Cre* | (107) | EMMA 00254<br>https://www.infrafrontier.eu/ | The Cre recombinase was cloned following an IRES sequence. An frt-flanked neomycin selection cassette was added, and the construct cloned in the 3' untranslated end of the tyrosine hydroxylase gene (as described by Althini et al., 2003, J Neurosci Res 72: 444-453). |

### Sex as a biological variable

Sex was not considered as a biological variable in these studies. Due to a previously reported increased variability of sodium appetite in female rodents (52), we performed all experiments in male mice.

### Stereotaxic injections

Mice were anesthetized with isoflurane 0.5–1.5% (titrated to respiratory depth and rate ∼40-80 breaths per minute with a deep surgical plane of anesthesia) then placed into a stereotaxic apparatus (Kopf 1900) with the head angled down. Meloxicam or carprofen (5 mg/kg s.c.) was provided for postoperative analgesia. After a midline incision, we retracted skin and muscle to expose the atlanto-occipital membrane atop the cisterna magna, then used a 28-gauge needle or scalpel to incise the membrane, providing direct access to the dorsal medulla. Through a pulled glass micropipette (20–30 µm tip inner diameter), we injected 90 nL of adeno-associated virus (AAV) targeting the caudal, medial NTS with coordinates 0.15 mm rostral, +/− 0.15 mm lateral, and 0.4–0.6 mm deep to calamus scriptorius. Each injection was made in 30 nL increments, 2 minutes each, at depths of 0.6, then 0.5, and finally 0.4 mm deep to calamus scriptorius, using picoliter air puffs through a solenoid valve (Clippard EV 24VDC) pulsed by a Grass stimulator. The pipette was left in place for an additional 5 minutes, then withdrawn slowly. The skin was then closed with Vetbond (3M).

To selectively ablate neurons that express Cre, we used AAV-DIO-dtA-mCherry (AAV1-Ef1a-Lox-Cherry-lox(dtA)-lox2.ape, 5.2 x 10^12^ vg/mL, purchased from Gene Therapy Center Vector Core, The University of North Carolina at Chapel Hill). To chemogenetically stimulate HSD2 neurons with i.p. injection, we used AAV-DIO-hM3Dq-mCherry (AAV1-Ef1a-DIO-hM3Dq-mCherry, 4 x 10^12^ vg/mL, purchased from the Duke viral vector core). In mice used for chemogenetically stimulating HSD2 neurons by CNO continuous infusion pumps, we used AAV8-DIO-hM3Dq-mCherry (Addgene #44361-AAV8, 2.1 x 10^13^ gc/mL), and control mice in this experiment received AAV8-DIO-mCherry (Addgene #50459-AAV8, 2.2 x 10^13^ gc/mL). After *post hoc* histology (described below in the “**Immunofluorescence labeling”** subsection), we included all experimental-ablation mice for analysis if they had fewer than 50% of the average number of HSD2 neurons counted control mice.

### Cannula implantation and subcutaneous osmotic minipumps

To infuse aldosterone into the fourth ventricle (i4V) or lateral ventricle (LV), we performed the following surgeries. Under anesthesia, the scalp was shaved, and the mouse placed gently in the stereotaxic ear-bars. A scalp incision was made wide enough to expose the skull at both bregma and lambda. After leveling the skull (both AP and ML) and zeroing at bregma, a 1 mm burr hole was made using a high-speed rotary tool with a 0.8 mm bit. An L-shaped 28-gauge cannula (3280PM-SPC, 5 mm; Plastics One) was attached by tubing (0.69 mm I.D. x 1.14 mm O.D., purchased from Scientific Commodities INC) to an osmotic minipump (ALZET 1007D) and then inserted through the burr hole to the target coordinate (i4V: 0 mm lateral, −6.5 mm caudal, and 5.0 mm deep to bregma for i4V; LV: 0.8 mm right, 0.35 mm caudal, and 2.5 mm deep to bregma). Cyanoacrylate glue with dental cement was used to fix the cannula. After the cannula was firmly cemented in place (about 4 minutes), the osmotic minipump was inserted into a subcutaneous pocket in the mid-scapular area. To infuse aldosterone peripherally, we placed the osmotic minipump in the same subcutaneous position without connecting any cannula or tubing. The skin incision was closed using Vetbond (3M).

At the end of each experiment in every mouse with an i4V or LV cannula, under anesthesia and immediately before transcardial perfusion (detailed below in the “**Transcardial perfusion and tissue sectioning**” subsection), we disconnected the catheter tubing from the osmotic minipump and used a 28-gauge insulin syringe to inject Evans blue dye (1% in sterile water) into the tubing to check patency of the cannula. This allowed us to determine, first, whether the dye entered without resistance, and then, via gross anatomical inspection and subsequent histological analysis, whether the dye flowed through the ventricular system and emerged into the basal cisterns and other subarachnoid spaces. This patency-check was particularly important for i4V cannula placements, due to blockage of the cannula by the choroid plexus or a gliotic membrane covering the cannula and redirecting fluid dorsally through the cerebellum, away from the fourth ventricle, in many cases. We excluded from analysis all mice that did not have Evans blue dye appear in the fourth ventricle or LV.

### BioDAQ fluid intake measurement

Mice were housed individually in BioDAQ cages (Research Diets), which allow precise fluid intake monitoring. All cages were housed on a stable shelf rack (MetroMax) to minimize vibration. Each BioDAQ cage has two highly precise, external scales (+/− 10 mg) continuously weighing a bottle of either water or saline. We did not vary placement of the water bottle (right) and 3% NaCl bottle (left) because appetite, not gustatory discrimination, was the subject of this study. We used 3% (0.51 M) NaCl, which provides optimal signal-to-noise for assessing sodium appetite; mice and rats have a baseline aversion to this saline concentration and typically drink zero to less than 0.1 mL per day unless sodium appetite is provoked (36, 53, 54). At the start of every experiment, each BioDAQ cage also had fresh ALPHA-dri+ Plus bedding (Shepherd Specialty Papers), an overhead wire hopper with standard rodent chow (Teklad 7013), and two bottles (distilled water and 3% NaCl) with sipper tubes, accessible through vestibules in the cage wall, that can be blocked by a gate. Between cohorts, we cleaned all cages, stainless steel cage mounts, liquid hoppers, blockers, and couplings. We also re-calibrated all scales at every cage-change, as recommended by the manufacturer, and all bottles were cleaned, then rinsed extensively with distilled water and refilled, between every cohort. Individual bottles were numbered and used in the same cage position, atop the same scale, for every cohort. Fluid intake was recorded continuously throughout every experiment, but we specified set intervals for analysis *a priori*, as detailed for individual experiments.

### Aldosterone dose-response testing

We infused aldosterone using ALZET 1007D osmotic minipumps (described above), which deliver 0.5 µL per hour from a 100 µL fluid reservoir. The day before implanting minipumps, we pre-loaded each pump with 100 µL of either vehicle (1% ethanol in sterile 0.9% saline) or an aldosterone solution, then placed the pump in 0.9% sterile saline overnight. We tested separate dose-ranges for i4V and subcutaneous infusion. We tested each dose in a separate group of mice, and each mouse only received a single dose.

For direct brain infusion, we monitored saline and water intake for up to 10 days in separate groups of mice receiving either vehicle i4V (n=4) or aldosterone i4V, at rates of 1 (n=3), 2.5 (n=2), 5 (n=6), 10 (n=5), 25 (n=5), 50 (n=4), or 100 (n=5) ng/h. After identifying a peak effect at 10 ng/h aldosterone (i4V), we used an additional group of mice to infuse that dose into the lateral ventricle (n=7). For one mouse in the 1 ng/h i4V infusion group, saline and water data from the 3 days of recording (2 pre-surgery and 1 day-of surgery) were aberrant due to a loose hopper, and we discarded the data from those three days for that one mouse only.

For systemic infusion, we monitored saline and water intake for up to 10 days in separate groups of mice receiving either vehicle s.c. (n=4) or aldosterone s.c. at rates of 10 (n=2), 100 (n=3), 250 (n=3), 500 (n=5), 750 (n=3), or 1000 (n=6) ng/h.

### Cell-type specific neuronal stimulation and ablation

We habituated mixed-background mice in individual BioDAQ cages for at least three days with low-sodium chow and *ad libitum* access to water and 3% NaCl. We then assessed saline intake in an ethologically relevant, high-aldosterone paradigm by closing the access gate for 3% NaCl (within 30 min before lights-off) so that mice would be sodium-deprived for the following six days of continuous access to water and low-sodium chow. Six days later, to assess sodium appetite, we re-opened the saline access gate 10 minutes before lights-off and measured 3% NaCl and water intakes across the following 6 hours.

Separately, we used a chemogenetic approach in *Hsd11b2-Cre* mice after injecting AAV-DIO-hM3Dq-mCherry into the NTS (as described above, at least four weeks earlier) to selectively stimulate HSD2 neurons. After at least three days of habituation to individual BioDAQ cages, we injected mice 30 min prior to lights-off with either the hM3Dq agonist clozapine-N-oxide (CNO, 1 mg/kg in sterile 0.9% NaCl with 1% DMSO) or sterile 0.9 % NaCl as a control (0.1 mL per 10 g mouse). After at least three days between tests, we repeated the injection with the other solution (vehicle or CNO), such that every mouse received both injections. Following each injection, we analyzed water and 3% NaCl intake across the 6 hours after lights-off. In additional cohorts of *Hsd11b2*-Cre mice, we injected AAV-DIO-hM3Dq-mCherry or AAV-DIO-mCherry (as above). After 3 weeks, we habituated mice to individual BioDAQ cages for 5 days, then implanted an ALZET 1007D minipump s.c. (as above) and continued monitoring water and 3% NaCl intake for 7d. At the end of each experiment, we perfused each mouse for histological assessment of HSD2 neuron transduction.

Next, in separate cohorts, we tested if HSD2 neurons are necessary for aldosterone-induced saline intake. For these tests, we used *Hsd11b2-Cre* experimental mice (n=19 i4V aldosterone; n=12 s.c. aldosterone) and their Cre-negative littermates as control mice (n=10 i4V aldosterone; n=6 s.c. aldosterone). All mice received stereotaxic microinjections of AAV-DIO-dtA-mCherry, as described above. After waiting four weeks for neuronal ablation, each mouse was placed in a clean BioDAQ cage. After measuring baseline water and 3% NaCl intake for at least three full days, we implanted osmotic minipumps, as described above, and returned each mouse to its BioDAQ cage to monitor fluid intake for 9 more days.

### Dietary sodium deprivation and potassium supplementation

For dietary sodium deprivation, we placed n=9 mice in individual clean cages with *ad libitum* access to water and low-sodium chow (TD-130591, Teklad/Envigo). To prevent consumption of any excreted sodium, the mouse was provided with a fresh, clean cage every day. A parallel control group (n=5) was run through the same protocol, except that this group had *ad libitum* access to regular chow. After 6 days, all mice were perfused transcardially (described below in the “**Transcardial perfusion and tissue sectioning**” subsection), and immediately before perfusion, we collected CSF and then blood from every mouse (described below in the “**Measurements in blood plasma and cerebrospinal fluid**” subsection).

For dietary sodium deprivation plus potassium supplementation, we modeled our approach after a similar dietary protocol followed by acute infusion of potassium chloride (KCl) in human subjects (55). However, instead of intravenous infusion at the end of our protocol, we gave a bolus of potassium by gavage (3M KCl, 0.1 mL per 10 g), similar to previous rodent experiments (56). Also, after pilot experiments testing a range of concentrations, we found that mice show no aversion to 1.5% or less KCl in drinking water (they consumed this solution at the same daily volumes as dH_2_O). Thus, our protocol for dietary sodium deprivation plus potassium supplementation was *ad libitum* access to 1.5% KCl in the drinking water, plus the same low-sodium chow as above, and fresh cage changes. After 6 days, these mice (n=8) were perfused immediately following CSF and then blood collection.

### Metabolic cage studies

To verify whether increased water intake after peripheral (s.c.) aldosterone infusion is secondary to an increase in urinary fluid loss, we used four groups of mice. Two groups were treated with aldosterone (1000 ng/h s.c.) and two groups with vehicle infusion. Before osmotic minipump implantation, mice were habituated for 3d in individual metabolic cages with *ad libitum* food and water access (Ugo Basile, 41700). Urine was collected and water and food intake were measured every day for 13d. After s.c. minipump implantation, mice were returned to the metabolic cages for 10d. In two groups (one aldosterone and one vehicle, n=4 each), water was offered *ad libitum*. The other two groups (one aldosterone and one vehicle, n=4 each) received 5 mL water (slightly more than vehicle-infused mice drink each day and less than s.c. aldosterone-infused mice drank in our initial BioDAQ recordings), every day for 7d, then *ad libitum* water access for the final 3d. In addition to expected evaporative fluid loss, 2–3 urine samples had fecal contamination in two mice in the 5 mL-water-restricted experiment, requiring removal of 2 data points from animal 7863 (days 6 and 10) and 3 data points from animal 7864 (days 1, 5, and 6).

### Measurements in blood plasma and cerebrospinal fluid (CSF)

After six days of aldosterone infusion or after dietary sodium deprivation, we collected blood and CSF. Prior to blood and CSF collection, mice receiving peripheral, subcutaneous infusion (vehicle, 250, 500, and 1,000 ng/h) had *ad libitum* access to a regular-chow diet and water. Mice receiving i4V infusion (vehicle, 5 and 10 ng/h) were monitored in BioDAQ cages with *ad libitum* access to regular chow along with tubes of dH_2_O and 3% NaCl as above. Because the CSF collection precluded our previous blue-dye assay for i4V cannula patency in these mice, we used a function, behavioral assay for infusion cannula patency and only collected blood and CSF from mice consuming at least 1 mL of 3% NaCl per day by infusion d6, which was at the low end of our prior dose-response results with 5 and 10 ng/h i4V aldosterone infusion (above). Mice consuming little to no 3% NaCl (in the range of vehicle-infused mice) were presumed to have a non-patent cannula and euthanized without blood or CSF collection.

To collect CSF, we used a straight, pulled glass capillary pipette (80–100 µm tip inner diameter). We anesthetized each mouse and used a surgical exposure similar to the NTS approach described above. However, instead of incising the atlantooccipital membrane, we inserted the pipette slowly through the membrane until CSF flowed spontaneously from the cisterna magna and into the capillary pipette. The pipette was left in place for 10–30 min. After slowly removing the pipette from the cisterna magna, CSF was ejected from the pipette into a 200 µL tube by inserting a 22 ga blunt needle into the pipette then using an attached 10 mL syringe to push air into the pipette. CSF was stored in the 200 µL tube at −80°C until removal for subsequent assays. CSF volumes collected from individual mice ranged typically 5–10 µL, but in some mice as little as 3 or as much as 12 µL was collected.

Immediately after collecting CSF, we injected ketamine/xylazine for terminal blood collection and euthanasia (described below in the “**Transcardial perfusion and tissue sectioning**” subsection). After five minutes, the peritoneum was opened and the IVC was punctured with an insulin syringe/needle (28 ga needle; 1 mL syringe) to collect typically 0.5 mL up to 1 mL of blood directly from the IVC. After collecting blood, we opened the thorax and proceed to euthanasia by transcardial perfusion as described below. Blood was ejected from the insulin syringe into a microcentrifuge tube then spun in a centrifuge for 5 min at 6,000 rcf at 4°C. Serum was extracted using a pipettor and stored at −80°C.

We used an enzyme-linked immunosorbent assay (ELISA) to measure the aldosterone concentration (ref# IB79134, IBL) in mouse blood plasma and pooled CSF. After thawing and diluting samples (1:10 for aldosterone), we followed the standard instructions for each commercially available sandwich ELISA kit. In a subset of mice receiving s.c. infusion of aldosterone (1000 ng/h) or vehicle, we used an ion-sensitive electrode to measure serum sodium (Prolyte; Diamond Diagnostics). To measure plasma osmolality, we also used an osmometer. In an additional cohort of mice receiving s.c. infusion of aldosterone (1,000 ng/h) or vehicle, we provided *ad libitum* access to food and water (no saline), for 7d, then we collected blood from the mandibular vein and centrifuged, extracted, and stored serum at −80 °C. In these samples, we measured copeptin with an ELISA kit (cat# MBS160428; MyBioSource, USA) used in previous work in mice (57). We also measured blood glucose using a glucometer (FreeStyle Lite; Abbott).

### Transcardial perfusion and tissue sectioning

All mice with stereotaxic AAV injection or implanted cannulas and a subset of mice with s.c. infusion pumps were transcardially perfused as follows. First, each mouse was anesthetized with ketamine (150 mg/kg) and xylazine (15 mg/kg), dissolved in sterile 0.9% saline and injected i.p. It was then perfused transcardially with phosphate-buffered saline (PBS) prepared from 10X stock (P7059, Sigma), followed by 10% formalin-PBS (SF100-20, Fischer Scientific). After perfusion, we removed each brain and fixed it overnight in 10% formalin-PBS. We sectioned each brain into 40 µm-thick coronal slices using a freezing microtome and collected tissue sections into separate, 1-in-3 series. We stored all tissue sections in cryoprotectant solution at −30 °C until further processing.

### Immunohistochemistry

For human and porcine brain tissue sections, we used nickel-diaminobenzene (NiDAB) immunohistochemistry to label HSD2. We removed tissue sections from cryoprotectant, rinsed them in PBS, and selected 16–20 total sections from one 1-in-12 series spanning approximately 1 cm of the caudal medulla oblongata. This tissue series contained intermediate through caudal levels of the human nucleus of the solitary tract, as well as the cuneate, gracile, and spinal trigeminal nuclei, plus the caudal inferior olivary complex and pyramidal tracts back through the spinomedullary transition. We next incubated sections for 30 minutes in 3% hydrogen peroxide (H_2_O_2_, #H325-100, Fisher) in PBT (PBS with 0.25% Triton X-100; BP151-500, Fisher) to quench endogenous peroxidase activity. After three PBS washes, we loaded sections into a primary antibody solution. This solution included a rabbit anti-HSD2 polyclonal antiserum (**Table 2**) in PBT with 2% normal donkey serum (NDS, 017-000-121, Jackson ImmunoResearch) plus 0.05% sodium azide (14314, Alfa Aesar) as a preservative. Sections were incubated in this primary antibody solution overnight, at room temperature, on a tissue shaker. The next morning, after three PBS washes, we incubated sections for 2 hours in a secondary antibody solution with biotinylated donkey anti-rabbit (1:500; #711-065-152 Jackson ImmunoResearch) in PBT-NDS-azide. After 3 more PBS washes, sections were placed for 1 hour in avidin-biotin complex (Vectastain ABC kit PK-6100; Vector), washed three times in PBS, then incubated in NiDAB solution for 10 minutes. Our stock DAB solution was prepared by crushing 100 tablets of diaminobenzidine (#D-4418, Sigma) with a mortar and pestle, then adding the resulting powder to 200 mL deionized water (ddH2O) and filtering the resulting suspension. We added 1 mL of this filtered DAB stock solution per 6.5 mL PBS. After 10 minutes in this DAB solution, we added hydrogen peroxide (0.8 µL 30% H_2_O_2_ per 1 mL PBS-DAB) and swirled sections for 1–3 min until observing gray-black color change. After three final washes in PBS, we mounted sections onto glass slides (#2575-plus; Brain Research Laboratories). Slides were then air-dried, dehydrated in an ascending series of alcohols and xylenes, and coverslipped using Cytoseal-60.

**Table 2.** Antisera used in this study.

| Antigen | Immunogen Description | Source, host species, RRID | Concentration |
| --- | --- | --- | --- |
| 11-beta-hydroxysteroid dehydrogenase type 2 (HSD2) | Amino acids 261-405 from the C-terminus of human 11β-HSD2 | Santa Cruz, rabbit polyclonal, cat# sc_20176, RRID: AB_2233199 | 1:500 (human & porcine tissue)<br>1:700 (mouse tissue) |
| 11-beta-hydroxysteroid dehydrogenase type 2 (HSD2) | Fusion protein Ag5146 (amino acids 1–405 encoded by human *HSD11B2*; GenBank BC036780) | Protein Tech, rabbit polyclonal, cat# 14192-1-AP, RRID: AB_2119643 | 1:2,000 (mouse tissue) |
| Choline acetyltransferase (ChAT) | ChAT from human placenta | Millipore, goat polyclonal, cat# AB144P, RRID: AB_2079751 | 1:500 (human tissue) |
| Fos | Synthetic peptide corresponding to residues near the amino terminus of rat c-Fos | Synaptic System, guinea pig; cat#226 308, RRID: AB_2905595 | 1:12,000 (mouse tissue) |
| mCherry | Full length mCherry fluorescent protein | Life Sciences, rat monoclonal, cat# M11217, RRID: AB_2536611 | 1:2,000 (mouse tissue) |
| Tyrosine hydroxylase (TH) | Native tyrosine hydroxylase from rat pheochromocytoma in sheep host | Millipore, sheep polyclonal, cat#: Ab1542, RRID: AB_90755 | 1:2,000 (mouse tissue) |

### RNAscope *in situ* hybridization

To label mRNA for human *HSD11B2*, we used the RNAscope 2.5 HD Detection Reagent-BROWN (ref#322310, lot#2006185; Advanced Cell Diagnostics, ACD) and a probe for human *HSD11B2* mRNA (cat# 432351, lot# 18165A; ACD). The afternoon before hybridization, we removed sections from cryoprotectant and rinsed them in PBS at room temperature before mounting them on glass slides kept at 4 °C. In the morning, we dehydrated the slides using 50%, 70%, and 2x 100% ethanol solutions for 5 min each. We then added 4 drops of H_2_O_2_ (RNAscope H_2_O_2_ and Protease Reagents, ref #322381, lot #2000897) to each slide for 10 minutes at room temperature. After two rinses with dH_2_O, we submerged slides into heated (100°C) Target Retrieval (RNAscope Target Retrieval Reagents, ref #322000,Lot# 2003928) for 15 min in a steamer (Oster). Next, the slides were submerged twice in dH_2_O for 2 min each, then quickly transferred to 100% ethanol for 1 minute. We used a Super-HI PAP pen (Research Products Incorporated) to form a hydrophobic barrier around the sections and added 4 drops of Protease plus. We then placed them on large glass petri dishes floating in a 40 °C water bath for 30 min. After 2 x 2 min washes in RNAscope wash buffer (ref 320058, lot# 2001175; diluted 1:50 in ddH2O) between each step, we incubated sections in the human *HSD11B2* probe (cat# 432351, lot# 18165A; ACD) for 2 hours at 40°C. After that, we added amplification reagents (Amp1-6) in series for 15-30 min each at 40°C, with 2 x 2 min washes in RNAscope wash buffer between each step. After Amp 6, we rinsed sections with wash buffer 2 x 2 min and added equal volumes of DAB-A and DAB-B for 10 minutes at room temperature. After a final wash with dH_2_O, we dehydrated slides through successive 1-minute dips in 50%, 70%, 2x 100% ethanol, and 2x xylene, then coverslipped each slide using Cytoseal-60.

### Immunofluorescence labeling

For immunofluorescence labeling of mouse brain tissue, we removed tissue sections from cryoprotectant and rinsed them in PBS before loading them into a solution containing one or more primary antisera (**Table 2**). These antisera were added to a PBS solution of 0.25% Triton X-100 (BP151-500, Fisher), 2% normal donkey serum (NDS, 017-000-121, Jackson ImmunoResearch), and 0.05% sodium azide (14314, Alfa Aesar) as a preservative (PBT-NDS-azide). We incubated sections overnight at room temperature on a tissue shaker. The following morning, the sections were washed 3x in PBS and incubated for 2 hours at room temperature in PBT-NDS-azide solution containing species-specific donkey secondary antibodies. These secondary antibodies were conjugated to Cy3, Cy5, Alexa Fluor 488, or biotin (Jackson ImmunoResearch #s 711-065-152, 711-165-152, 711-175-152, 705-065-147, 705-545-147, 713-545-147, 706-545-148, 706-165-148, 706-065-148, 715-065-15, 712-165-153; each diluted 1:1,000 or 1:500). If a biotinylated secondary antibody was used, sections were again washed 3x in PBS, then incubated for an additional 2 hours in streptavidin-Cy5 9#SA1011; Invitrogen) or streptavidin-Pacific Blue (#S11222, Life Technologies) diluted 1:1,000 in PBT-NDS-azide. Tissue sections were then washed 3x in PBS, mounted on glass slides (#2575-plus; Brain Research Laboratories), dried, and then coverslipped using Vectashield (Vector Labs). All slides were stored in slide folders at 4 °C until imaging.

### Imaging, cell counts, and figures

All slides were scanned using an Olympus VS120 microscope. Using the VS-ASW software (Olympus), we first acquired a 2x overview scan, then used an 10x objective to scan all tissue sections. Next, we acquired 20x and, in some cases, 40x z-stacks encompassing the NTS at every level containing HSD2 neurons. For each slide, this produced a virtual slide image (VSI) file containing a 2x and a 10x whole-slide layer, plus separate layers with 20x and/or 40x extended-focus images in regions of interest. We reviewed all slides in OlyVIA (Olympus) and used VS-ASW or cellSens (Olympus) to crop full-resolution images, and we used Adobe Photoshop to adjust brightness and contrast. We used Adobe Illustrator to arrange panels and add lettering for final figure layouts. Scale bars were traced atop calibrated lines from cellSens, in Illustrator, to produce clean white or black lines.

For brightfield analysis in human tissue, we reviewed all slides at high magnification to identify HSD2-immunoreactive and neuromelanin-containing neurons throughout each tissue section. After selecting three adult brainstems with the highest tissue quality (MH001, MH004, and MH005), we used VS-ASW to count every HSD2-immunoreactive neuron that contained a nuclear void and measured the short-axis diameter of each neuron through the center of its nucleus (in µm). We then used the average cell diameter to perform Abercrombie correction (58), multiplied this number by the number of tissue series (x12 for 1-in-12 series) to estimate the total number of HSD2 neurons in that case, and averaged this estimate across cases.

For epifluorescence analysis in mouse tissue, we used QuPath (59) to count all NTS neurons that were immunoreactive for HSD2, as well as all neighboring, catecholaminergic neurons in the NTS that were immunoreactive for the enzyme tyrosine hydroxylase (TH). Mice with tears in the dorsal hindbrain that sometimes occurred during brainstem removal, due to post-surgical meningeal adhesions, were excluded from analysis if the resulting histological artifacts prevented analysis of the full caudal NTS (n=5 cases from *Th*-IRES-Cre ablation and control groups).

### Data availability

The data that support the findings of this study are available from the corresponding author upon reasonable request.

### Data analysis and statistics

We exported continuous fluid intake records from the BioDAQ DataViewer (Research Diets), then used Microsoft Excel to organize the data and to calculate total intake volumes of water and 3% NaCl. After acute chemogenetic stimulation with CNO and after week-long dietary sodium deprivation, we analyzed the first 6 hours of access to 3% NaCl, as in previous work (54). For all multi-day infusion protocols, we analyzed 3% NaCl and water intake in 24-hour bins. We then used GraphPad Prism to plot data and run statistical tests. To compare dose-response effects of aldosterone on 3% NaCl and water intake, we calculated area under the curve (AUC) across 9d following osmotic minipump implantation, then used one-way ANOVA followed by Tukey’s multiple comparisons correction. To assess the effect of Cre-conditional neuronal ablation on aldosterone-induced intake of 3% NaCl and water, CNO effects on sodium and water intake, and sodium and water intake induced by sodium deprivation, we used unpaired, two-tailed t-tests. We also used unpaired, two-tailed t-tests to compare immunolabeled cell counts between individual groups of experimental and control mice and to compare sodium, copeptin, and glucose measurements between aldosterone-infused and control mice. To compare average daily water intake and average daily urine output between aldosterone- and vehicle-infused mice with restricted and unrestricted water access in metabolic cages, we used repeated measures two-way ANOVA. All results are expressed as mean ± standard deviation.

### Study approval

All procedures involving animals were conducted in accordance with the guidelines of the Institutional Animal Care and Use Committees at the University of Iowa (protocol numbers 00720011, 3072011, and 3102343).

## Results

### Human HSD2 neurons

Some neural circuits are different between rodents and humans, and our interest in this topic extends only so far as it is relevant to human health. Therefore, we began by testing whether there are HSD2 neurons in the human brain. In the caudal nucleus of the solitary tract (NTS), we found immunohistochemical labeling for HSD2 protein (N=6) and *in situ* hybridization for *HSD11B2* mRNA (N=3). As shown in **Figure 1A–C** (and in **Supplemental Figure 1D**), the location, morphology, and distribution of these neurons in the human NTS closely resembles the HSD2 population we identified in rodents (29, 30). As in rats and mice, human HSD2 neurons (red dots in **Figure 1D**) lie near the obex of the fourth ventricle and extend caudally, through the commissural NTS, almost back to the spinomedullary transition. Also like HSD2 neurons in rats and mice, human HSD2 neurons neighbor a large population of neuromelanin-containing (catecholaminergic) neurons that extends laterally through the NTS and into the medullary reticular formation (green dots in **Figure 1D**). Human HSD2 neurons also neighbor a ventrolateral population of cholinergic neurons, in the dorsal vagal motor nucleus, which are immunoreactive for choline acetyltransferase (ChAT; **Figure 1F**).

**Figure 1.**
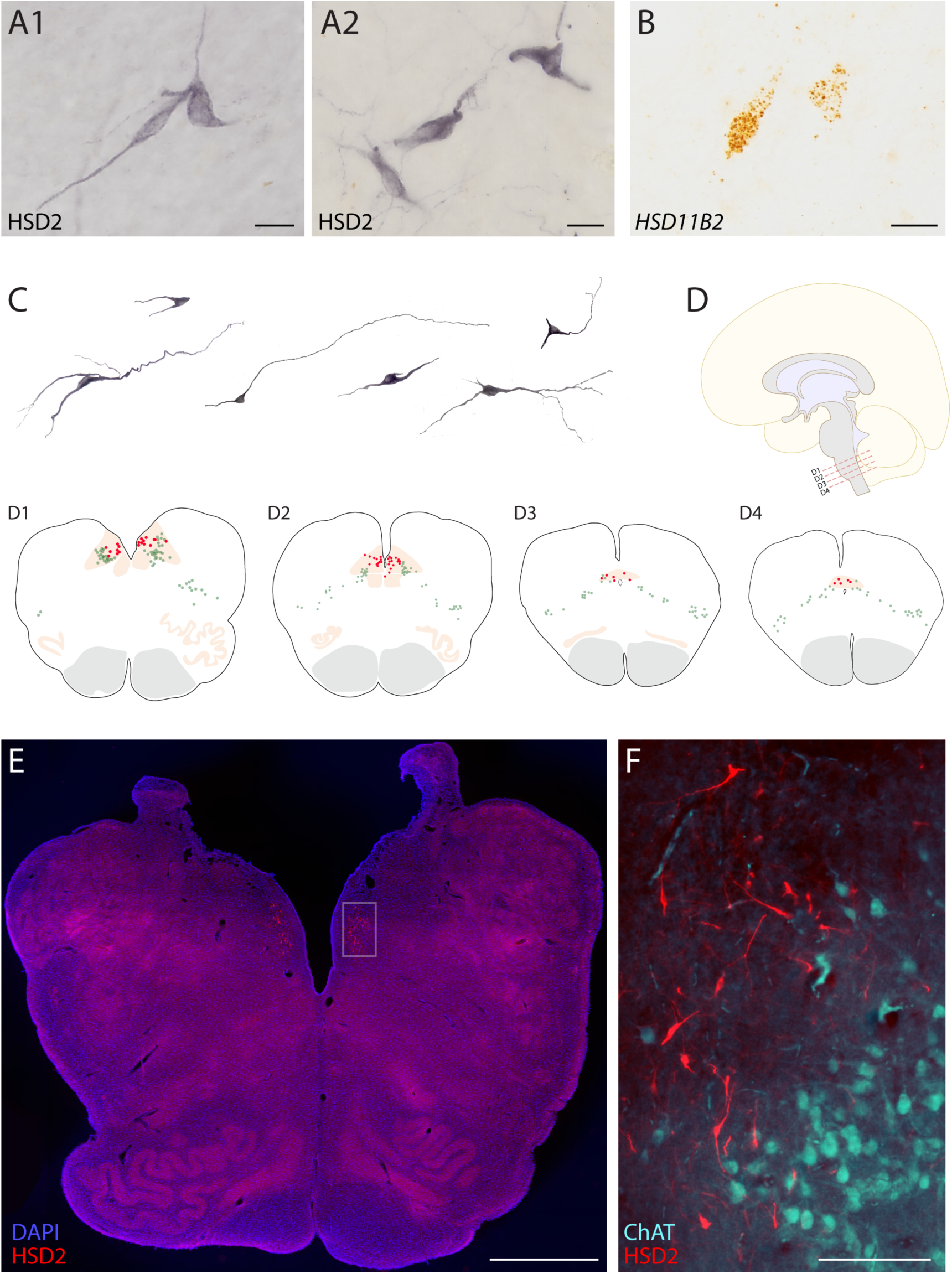
HSD2 (11-beta-hydroxysteroid dehydrogenase 2) neurons in the human brain. (A) Representative HSD2-immunoreactive neurons in the nucleus of the solitary tract (NTS) of two human cases (nickel-DAB immunohistochemical labeling from MH001 in A1 and MH005 in A2). (B) Representative *HSD11B2* mRNA labeling in the medial NTS (DAB *in situ* hybridization from case MH001). (C) Additional examples of HSD2 neuronal morphology across several human cases (left to right: MH005, MH001, MH001, MH004, MH001, and MH005). (D) Neuroanatomical location and distribution of HSD2 neurons in the human brain (red dots), along with neuromelanin-containing neurons (green dots) in the medulla oblongata (plots from four rostrocaudal levels of the caudal brainstem in case MH005). HSD2-immunoreactive neurons are restricted to the caudal-medial NTS, and their distribution runs from the obex of the fourth ventricle (D1–D2) back through the caudal commissural NTS (D3–D4), just before the spinomedullary transition. (E) HSD2 immunofluorescence in NTS neurons flanking the caudal fourth ventricle. (F) Combined HSD2 (red) and choline acetyltransferase (ChAT, ice-blue) immunofluorescence labeling reveals the distinction between HSD2 neurons in the medial NTS and cholinergic neurons in the dorsal motor nucleus of the vagus nerve. Scale bars in (A–B) are 20 µm. Scale bars in (E) and (F) are 2 mm and 200 µm, respectively.

Human HSD2 neurons are spindle-shaped, with two or three primary dendrites. The short-axis diameter of each HSD2 neuronal soma averaged 10.9 ± 2.5 µm (range 2.5–18.9 µm; n=278 neurons from N=3 human brainstems). From rostrocaudal, Abercrombie-corrected counts of HSD2 neurons from three cases, we estimate that the human brainstem contains 958 ± 320 HSD2 neurons (1252 in #MH001, 617 in #MH004, 1003 in #MH005), roughly twice the number of HSD2 neurons in the rat brainstem (29).

In addition to rodents and humans, we also detected HSD2-immunoreactive neurons in pigs, in the medial NTS, near the obex of the fourth ventricle (**Supplemental Figure 1C**). Thus, HSD2 neurons have a similar location, distribution, and morphology in several mammalian species, at least including rodents (mouse and rat), ungulates (pig), and primates (human).

### Sodium deprivation activates HSD2 neurons and their chemogenetic stimulation increases saline intake

Switching rats to a low-sodium diet for one week is a simple and ethologically relevant way to boost aldosterone production, activate HSD2 neurons, and increase sodium appetite (34, 54, 60). These effects are uncharacterized in mice, so we first assessed the impact of sodium deprivation on HSD2 neurons and sodium appetite. Switching mice to a low-sodium chow (<0.01% Na) increased the proportion of HSD2 neurons expressing Fos, a neuronal activity marker (**Figure 2A-B**), and a parallel group consumed more saline when given access to a 3% NaCl drinking tube after 6d dietary sodium deprivation (**Figure 2C**). Thus, as in rats, dietary sodium deprivation activates HSD2 neurons and increases sodium appetite in mice.

**Figure 2.**
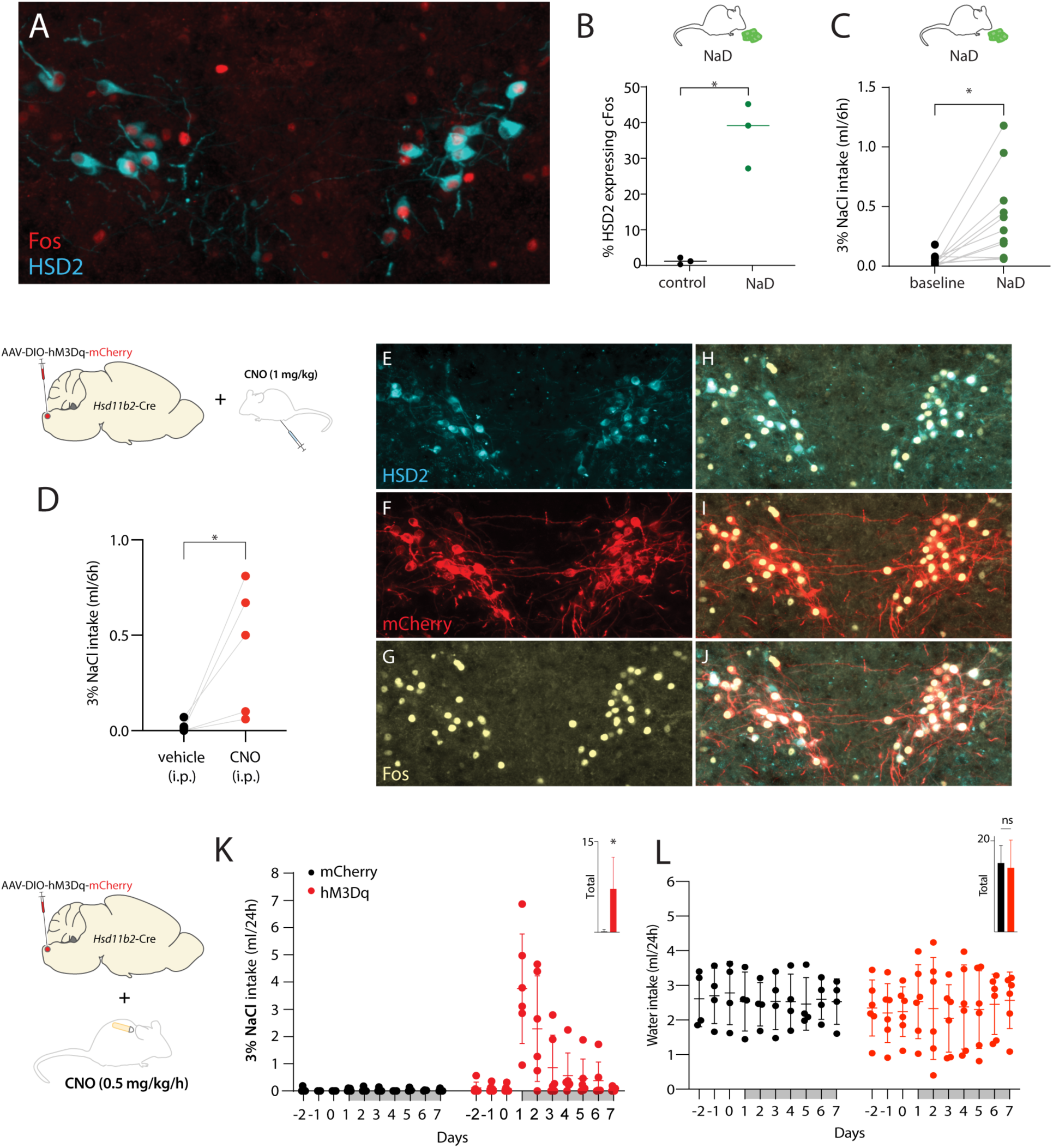
Effects of dietary sodium deprivation and chemogenetic stimulation on HSD2 neuron activity and salt consumption. (A–B) Feeding mice low-sodium chow (<0.01% Na) for 6 days increased the percentage of Fos-immunoreactive HSD2 neurons (p=0.0025, versus control mice with *ad libitum* access to 0.3% Na regular chow). (C) Dietary sodium deprivation also caused mice to consume more saline (p=0.0042 versus intake of 3% NaCl during access to regular, 0.3% Na chow). (D) Acute chemogenetic activation of HSD2 neurons by injection of clozapine-N-oxide (CNO, 1 mg/kg i.p.) increased saline intake (p=0.0490 versus vehicle injection). (E–J) CNO (1 mg/kg, 60 min prior to perfusion) caused Fos activation of HSD2 neurons expressing Cre-conditional hM3Dq-mCherry. (K–L) Using an osmotic minipump to continuously infuse CNO (0.5 mg/kg/h) produced a large increase in saline intake (p=0.0297) without changing water intake (p=0.7675).

Next, we extended previous reports that chemogenetically stimulating HSD2 neurons increases sodium appetite (35, 36). First, we used *Hsd11b2*-Cre mice (n=5) to selectively express the excitatory DREADD (designer receptor exclusively activated by designer drug) hM3Dq-mCherry in HSD2 neurons. In these mice, we confirmed that injecting the hM3Dq ligand clozapine-N-oxide (CNO, 1 mg/kg i.p.) robustly activates HSD2 neurons (**Figure 2E-J**) and increases saline consumption (**Figure 2D**). In additional groups, we implanted osmotic minipumps to infuse CNO (0.5 mg/kg/h) in mice with either hM3Dq-mCherry or mCherry expressed selectively in HSD2 neurons. Continuous CNO infusion increased 3% NaCl intake to large daily volumes (**Figure 2K**), higher than any amount of sodium intake reported previously in mice. This large natriorexigenic effect of continuous CNO infusion was mediated by hM3Dq-mCherry because CNO had no effect in mCherry control mice. Continuous activation of HSD2 neurons selectively increased saline intake (p=0.0297) without altering water intake (p = 0.7675; **Figure 2L**).

### Aldosterone dose-dependently increases saline intake

Previous studies in mice reported that mineralocorticosteroids do not increase sodium appetite (37, 38), but none of these studies tested aldosterone, which is the primary mineralocorticosteroid hormone. Infusing aldosterone directly into the fourth ventricle (i4V) produced voracious saline intake in rats (11, 12), so we tested whether i4V aldosterone infusion can increase sodium appetite in mice. We began with the infusion rate that had increased saline intake in rats (100 ng/h), but this dose did not increase saline intake, and 3 of 5 mice either died unexpectedly or had to be euthanized due to low fluid intake. Due to this unexpected lethality at 100 ng/h, we performed a series of dose-ranging experiments at lower infusion rates (1–50 ng/h) and recorded 3% NaCl and water intake continuously, in individually housed mice, for 2d prior through 9d after implanting an osmotic minipump infusing aldosterone through an i4V cannula. We verified cannula patency *post hoc* by infusing dye and included mice for analysis only if the dye passed into the fourth ventricle without resistance (**Figure 3A**). Our primary endpoint was total (9d) 3% NaCl consumption. We also examined total water consumption and time courses of saline and water intake.

**Figure 3.**
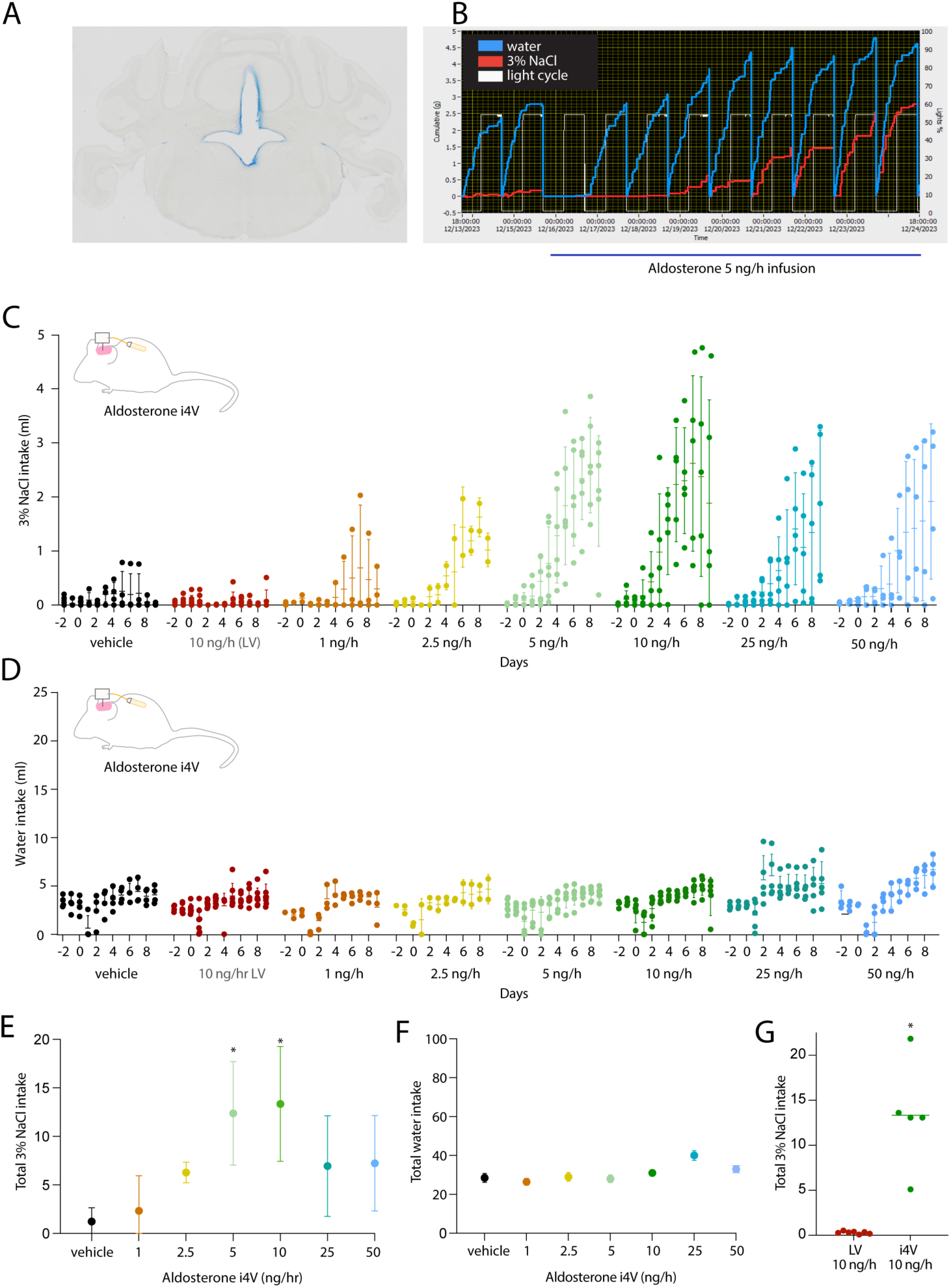
Dose-response relationship between fourth cerebral ventricle (i4V) infusion of aldosterone and 3% NaCl intake. (A) Cannula patency was tested by injecting Evan’s blue dye at the end of the experiment. In all mice included for analysis, dye passed through the cannula, without resistance, and appeared in the fourth ventricle. (B) Water (blue) and 3% NaCl (red) BioDAQ intake data for a representative mouse with i4V aldosterone infusion (5 ng/h). The recording begins 2d before and continues for 9d after cannula and minipump implant. (C) 3% NaCl intake of i4V aldosterone- and vehicle-infused mice, plus one group with aldosterone 10 ng/h infusion into the lateral ventricle (LV, red). In addition to individual mouse data, each group mean (horizontal line) and standard deviation (vertical line) are shown for each day. (D) Water intake of i4V aldosterone- and vehicle-infused mice, plus one group with aldosterone 10 ng/h infusion into the LV (red). (E-F) Total (9d) 3% NaCl (E) and water (F) intake across i4V groups. (G) Total (9d) 3% NaCl intake induced by LV infusion (red) and by i4V infusion (green) of 10 ng/h aldosterone. Asterisks indicate a statistically significant difference (p<0.05 by one-way ANOVA followed by Tukey’s test for multiple comparisons: p=0.020 for 3% NaCl in the 5 ng/h group; p=0.0139 for 3% NaCl in the 10 ng/h group).

Vehicle infusion had no effect, but i4V aldosterone infusion caused a prominent, dose-dependent rise in saline consumption. **Figure 3B** shows the raw intake data before and during infusion of 5 ng/h aldosterone in a representative mouse (3% NaCl in red; water in blue). Saline intake increased gradually to a plateau by day 7, and we found an inverted U-shaped relationship between i4V aldosterone dose and saline intake behavior, with peak effects at 5–10 ng/h and reduced potency both above and below this range (**Figure 3E**). Mice receiving 5 or 10 ng/h i4V aldosterone consumed the most saline, and both groups consumed more 3% NaCl than vehicle-infused mice, without a statistically significant difference between these two doses (**Figure 3E**). The appetitive state induced by i4V aldosterone infusion was selective for saline; some mice drank slightly more water, but there was no statistically significant effect at any i4V aldosterone infusion rate, relative to vehicle (**Figure 3F**). Infusing a maximally effective i4V dose (10 ng/h) into the lateral ventricle had no effect on saline intake (p=0.0001; **Figure 3G**) or water intake (p=0.135). The potent effect of aldosterone in the fourth ventricle, but in not the lateral ventricle, points to a site of action in the hindbrain rather than the forebrain.

We next characterized the effect of peripheral aldosterone infusion. We tested a range of infusion rates to reproduce plasma aldosterone levels spanning the full range of aldosterone levels reported after dietary sodium deprivation and across a variety of pathophysiologic states in humans, including dietary sodium deprivation, hyperkalemia, kidney disease, and aldosterone-secreting adenomas (34, 55, 61, 62). We again found dose-dependent effects on saline intake. However, the i4V maximally effective infusion rate of 10 ng/h had no effect (**Figure 4A**), and peripheral aldosterone infusion was 100-fold less potent than i4V infusion (**Figure 4C**). Unlike the inverted U-shaped response to i4V infusion, we found a more linear effect, with the largest peripheral infusion rate (1,000 ng/h) producing the maximum behavioral response. Infusing 750 ng/h had a smaller effect (p=0.0040), but both 750 ng/h and 1,000 ng/h aldosterone infusion induced significantly more saline intake than vehicle (p=0.0119 and p<0.0001). In contrast to the salt-specificity of i4V infusion, peripheral aldosterone infusion also increased water intake (**Figure 4B,D**). Elevated water intake was not evident in mice receiving vehicle or 10 ng/h aldosterone, but an effect on water intake was apparent at all peripheral infusion rates above 10 ng/h. Mice receiving 100, 250, and 500 ng/h increased their water intake approximately three-fold, and those receiving 750 and 1,000 ng/h increased nearly five-fold. Plotting total saline intake versus water intake for every mouse (**Figure 4E**) confirmed that i4V infusion selectively increased saline intake, while peripheral infusion increased both saline and water intake.

**Figure 4.**
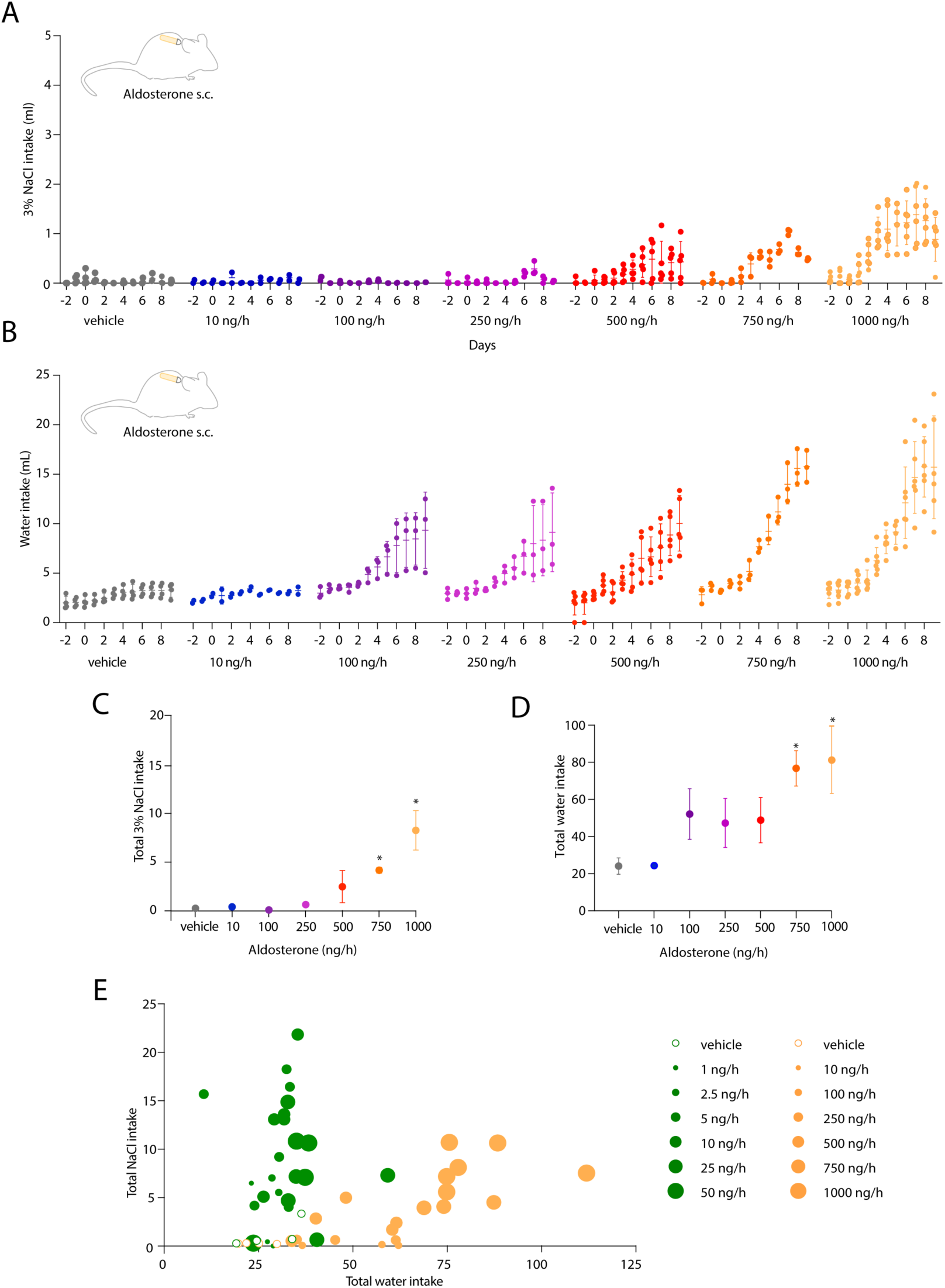
Dose-response relationship between subcutaneous (s.c.) aldosterone infusion and 3% NaCl intake. (A) 3% NaCl intake of aldosterone- and vehicle-infused mice. Group mean (horizontal line) and standard deviation (vertical line) are shown for each day, along with individual mouse data. (B) Water intake of aldosterone- and vehicle-infused mice. (C-D) Total (9d) 3% NaCl (C) and water (D) intake across all s.c. infusion groups. Asterisks indicate a statistically significant difference by one-way ANOVA followed by Tukey’s test for multiple comparisons (p=0.0119 for 3% NaCl in the 750 ng/h group; p<0.0001 for 3% NaCl in the 1000 ng/h group; p=0.0043 for water in the 750 ng/h group; p=0.0006 for water in the 1000 ng/h group). (E) Scatterplot showing total saline intake (y-axis) versus total water intake (x-axis) for every mouse. Dot size indicates rate and color indicates route of infusion (i4V green; s.c. orange), as shown in the legend.

The extent to which aldosterone crosses the blood-brain barrier remains unclear (47, 63), leaving open the possibility that i4V infusions that increased saline intake did so by producing aldosterone concentrations that are supraphysiologic in brain tissue. To address this possibility, we measured endogenous aldosterone concentrations in a variety of conditions, in both blood and cerebrospinal fluid (CSF), then assessed whether our i4V infusions produced CSF concentrations within or outside the endogenous range. We also tested whether and to what extent peripheral aldosterone infusions increase the aldosterone concentration in CSF. As in rats (34), removing dietary sodium increased aldosterone production, with blood plasma concentrations reaching 113 to 414 ng/dL (**Figure 5A**). These sodium-deprived mice also had elevated aldosterone concentrations in CSF, ranging 19 to 47 ng/dL (**Figure 5B**). To determine the high end of endogenous production, we adapted a human protocol that combined dietary sodium deprivation with potassium supplementation, followed by acute potassium infusion (55). Supplementing dietary potassium in sodium-deprived mice, followed by potassium gavage (56), produced higher concentrations of endogenous aldosterone in blood plasma (ranging 313 to 990 ng/dL, **Figure 5A**) and in CSF (ranging 48 to 74 ng/dL, **Figure 5B**).

**Figure 5.**
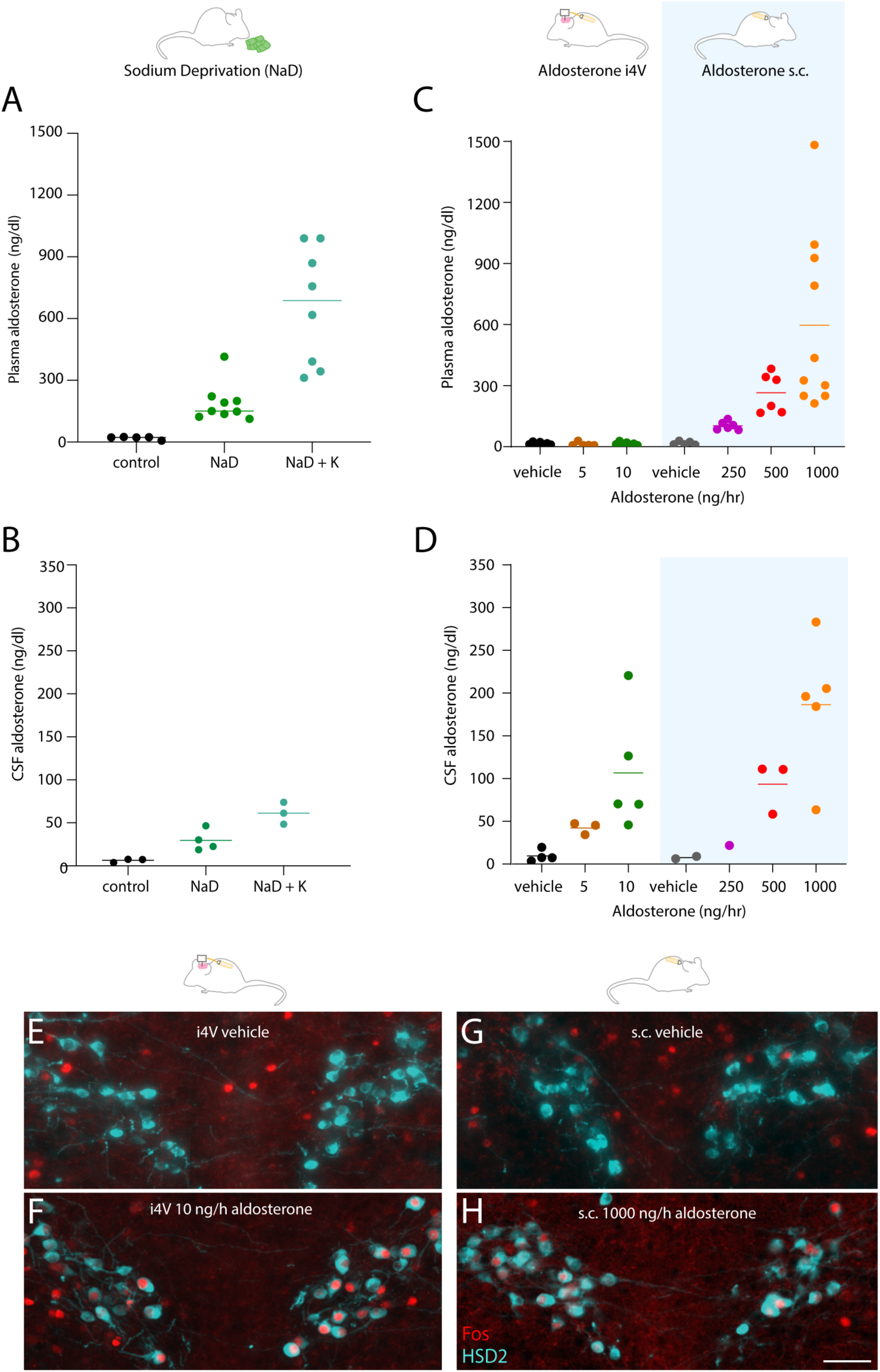
Effects of diet and aldosterone infusion on the aldosterone concentration in blood plasma and cerebrospinal fluid (CSF). (A) Plasma aldosterone concentrations of mice after dietary sodium deprivation (NaD) and after NaD with potassium supplementation (NaD+K). (B) CSF aldosterone concentrations of mice after NaD and NaD+K. Each dot represents pooled CSF from 2–3 mice. (C) Plasma aldosterone concentrations of mice receiving aldosterone i4V infusion (5–10 ng/h) or s.c. infusion (250– 1000 ng/h). (D) CSF aldosterone concentrations of mice receiving i4V and s.c. infusions. Each dot represents pooled CSF from 2–3 mice. For each group with more than one sample, the mean is represented by a small horizontal line with the same color as individual samples in that group. (E–H) Aldosterone i4V (E) and s.c. (G) induced Fos immunoreactivity in HSD2 neurons, and vehicle infusions did not (F,H).

After determining the physiologic ranges of endogenous aldosterone in blood and CSF, we next examined the ranges of aldosterone concentrations produced by aldosterone infusion. As expected, i4V infusion increased the aldosterone concentration in CSF (**Figure 5D**), with no apparent change in blood plasma (**Figure 5C**). In contrast, peripheral infusion increased the aldosterone concentration in both blood plasma and CSF (**Figure 5C-D**), indicating that some aldosterone had crossed the blood-brain barrier. Importantly, i4V infusion rates that had induced maximal salt intake did so at CSF aldosterone concentrations overlapping the range of endogenous aldosterone concentrations produced by one week of low-sodium diet. Specifically, mice receiving 5 ng/h i4V infusion had CSF aldosterone concentrations ranging 34 to 47 ng/dL (**Figure 5D**), which closely resembled CSF aldosterone concentrations after one week of dietary sodium deprivation (19 to 47 ng/dL; **Figure 5B**). Altogether, our results indicate that circulating aldosterone reaches the brain and maximally increases salt intake at tissue concentrations between 34 to 47 ng/dL (0.9–1.3 nM). These concentrations are strikingly similar to aldosterone’s known functional range for binding and transcriptional activation of the mineralocorticoid receptor, which is approximately 0.1–1 nM (64–67).

### Aldosterone activates HSD2 neurons

Like other stimuli that increased saline intake (**Figure 2**), infusing aldosterone induced Fos expression in HSD2 neurons. We found substantial Fos activation of HSD2 neurons in mice receiving central (i4V, **Figure 5E**) and peripheral (s.c., **Figure 5G**) infusion.

### HSD2 neurons are required for aldosterone-induced saline intake

Defining a dose-response relationship between aldosterone and sodium intake allowed us to next focus on our main hypothesis, that HSD2 neurons are necessary for aldosterone-induced salt intake. To test this hypothesis, we infused aldosterone at maximally effective i4V and s.c. doses (10 ng/h and 1000 ng/h, respectively), in separate groups of experimental and control mice. In experimental mice, we selectively eliminated HSD2 neurons by delivering Cre-dependent diphtheria toxin A (AAV1-mCherry-DIO-dtA; **Figure 6B**) bilaterally into the NTS of *Hsd11b2*-Cre mice. As a control, we injected the same Cre-dependent vector into the NTS of Cre-negative littermates (**Figure 6A**). At the end of each experiment, we counted the number of remaining HSD2 neurons at every rostrocaudal level of the NTS and found that the Cre-dependent ablation approach substantially reduced the number of HSD2 neurons (p<0.0001 relative to Cre-negative littermates; **Figure 6C**).

**Figure 6.**
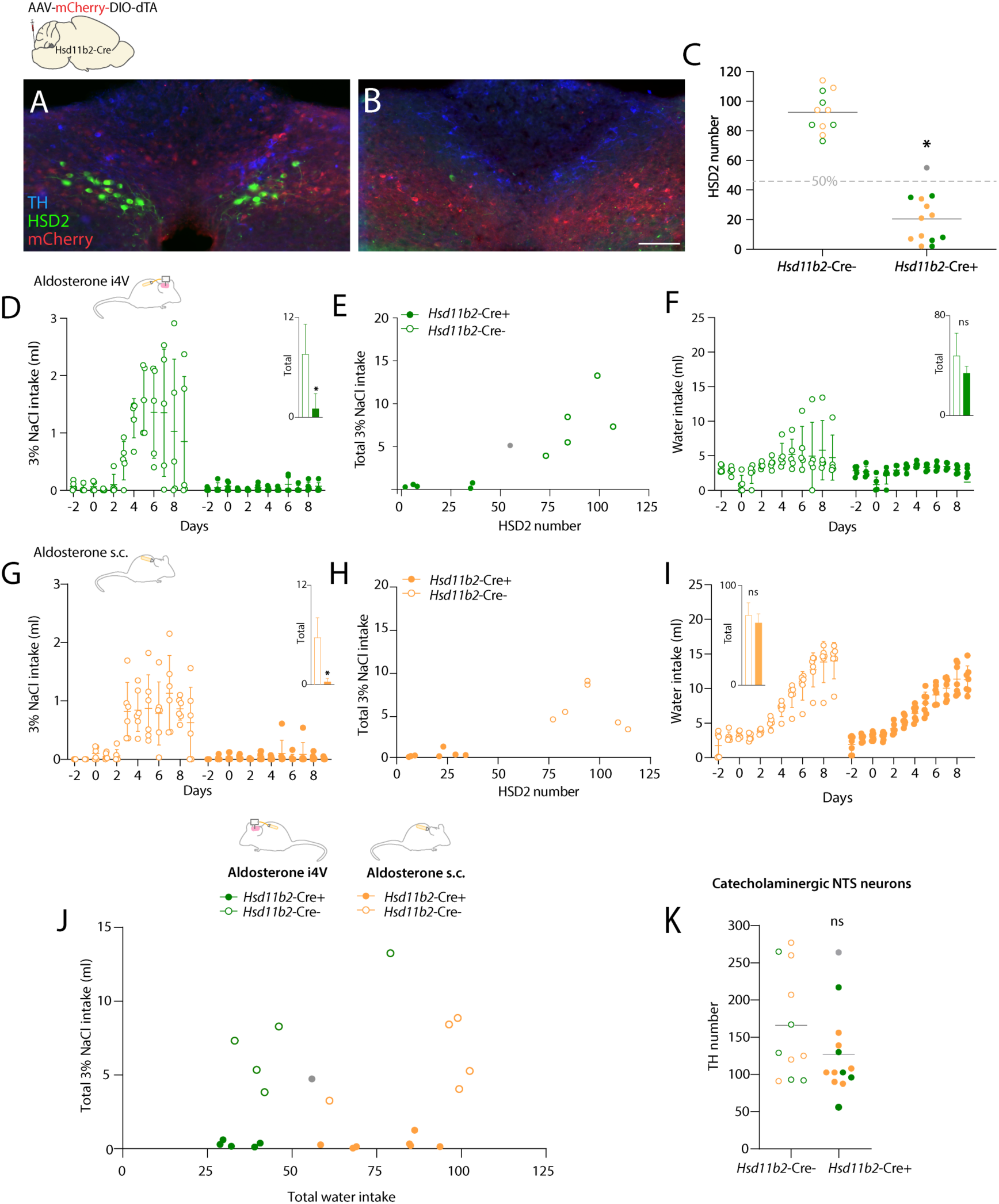
Effect of HSD2 neuron ablation on aldosterone-induced salt intake. (A) After bilateral microinjections of the ablation vector (AAV-mCherry-DIO-dtA) into the nucleus of the solitary tract (NTS), control mice (Cre-negative littermates) had a normal distribution of neurons that are immunoreactive for 11-beta-hydroxysteroid dehydrogenase (HSD2, green). With this AAV, non-Cre expressing neurons in the injected region express mCherry. (B) Experimental mice (*Hsd11b2*-Cre+) had few remaining HSD2 neurons. In both (A) and (B), the neighboring population of catecholaminergic NTS neurons are evident by their immunoreactivity for tyrosine hydroxylase (TH, blue). (C) Genetically targeted ablation successfully reduced the number of HSD2 neurons (*p<0.0001 by two-tailed t-test). One case (#3383) is represented by a gray dot, signifying exclusion from subsequent analysis because it did not meet the prespecified criterion of eliminating at least 50% of the average HSD2 neuron number in control mice. (D–F) Effects of HSD2 neuronal ablation on 3% NaCl and water intake in mice receiving a maximally effective rate of aldosterone infusion (10 ng/h i4V). The gray dot in (E) again represents case #3383, which did not meet the prespecified criterion of >50% ablation. In panel (D) inset, * indicates p=0.0034. In panel (F) inset, “ns” indicates “not significant” (p=0.1368). (G-I) Effects of HSD2 neuron ablation on 3% NaCl and water intake in mice receiving a maximally effective rate of s.c. aldosterone infusion (1000 ng/h). In panel (G) inset, * indicates p<0.0001. In panel (I) inset, “ns” indicates “not significant” (p=0.2106). (J) Scatterplot showing total saline intake (y-axis) versus total water intake (x-axis) for every mouse (i4V green; s.c. orange; Cre-negative littermate control mice, open circles; *Hsd11b2*-Cre+ ablation mice filled circles). (K) Genetically targeted ablations in *Hsd11b2*-Cre+ mice did not alter the number of neurons in the neighboring population of NTS catecholaminergic (TH-immunoreactive) neurons (p=0.1584).

In Cre-negative control mice, as in the dose-response tests above, infusing aldosterone robustly increased saline intake. In contrast, ablating HSD2 neurons prevented saline intake induced by i4V aldosterone infusion (p=0.0034 relative to Cre-negative littermates, **Figure 6D-E**). In separate groups of ablation and control mice, ablating HSD2 neurons also prevented saline intake induced by s.c. aldosterone infusion (p<0.0001 relative to Cre-negative littermates, **Figure 6G-H**).

Examining the relationship between saline intake and number of remaining HSD2 neurons (scatterplots in **Figure 6E,H**) suggests that aldosterone-induced sodium appetite can be prevented by eliminating 50% of the HSD2 neuron population, because mice with up to 36 (i4V) or 34 (s.c.) remaining HSD2 neurons consumed little to no saline during aldosterone infusion. Only one *Hsd11b2*-Cre+ mouse failed our inclusion criterion of >50% ablation (gray dot in **Figure 6C** and **6E**), and this mouse – with 55 surviving HSD2 neurons and a total (9d) saline intake of 5.13 mL – was the only mouse in the experimental group with an aldosterone-induced salt intake in the range of control mice.

We also examined whether eliminating HSD2 neurons alters water intake. As in our initial dose-response tests, i4V infusion selectively increased saline intake in control mice, with little to no change in water intake, while s.c. infusion increased both saline and water intake in control mice (**Figure 6G,I**). In contrast to the total loss of aldosterone-induced saline intake, the elevated water intake caused by peripheral (s.c.) aldosterone remained prominent in *Hsd11b2*-Cre+ experimental mice (**Figure 6I**), indicating that HSD2 neurons selectively mediate aldosterone-induced sodium appetite, not thirst.

To better assess the ablation specificity in every mouse, we also counted the number of remaining neurons in the large, neighboring population of NTS catecholaminergic neurons. These neurons are immunoreactive for the enzyme tyrosine hydroxylase (TH), and none express HSD2 (29, 30, 68, 69). Cre-dependent ablations of HSD2 neurons in *Hsd11b2*-Cre mice did not alter the number of NTS catecholaminergic neurons (p=0.1584 relative to Cre-negative littermates; **Figure 6K**).

We next tested whether aldosterone-induced sodium appetite depends on HSD2 neurons exclusively, or on neighboring NTS neurons as well. We used a similar, genetically targeted approach to ablate NTS catecholaminergic neurons in *Th*-IRES-Cre mice (**Figure 7A-B**). Our approach selectively eliminated TH-immunoreactive neurons in *Th*-IRES-Cre+ mice (p=0.04 relative to Cre-negative littermates; **Figure 7C**), without altering the number of HSD2 neurons (p=0.2886; **Figure 7D**). All experimental mice (*Th*-IRES-Cre) and all control mice (Cre-negative littermates) exhibited aldosterone-induced saline intake (**Figure 7D**) with no difference between groups (p=0.7120; inset in **Figure 7D**). Saline consumption in these mice had no apparent relationship to the number of surviving catecholaminergic neurons in the NTS (**Figure 7E**). All mice also exhibited a prominent rise in water intake during s.c. aldosterone infusion, with no difference between groups (p=0.3315; **Figure 7F**). Thus, aldosterone-induced sodium appetite specifically requires HSD2 neurons, and not the larger, neighboring population of catecholaminergic neurons in the NTS.

**Figure 7.**
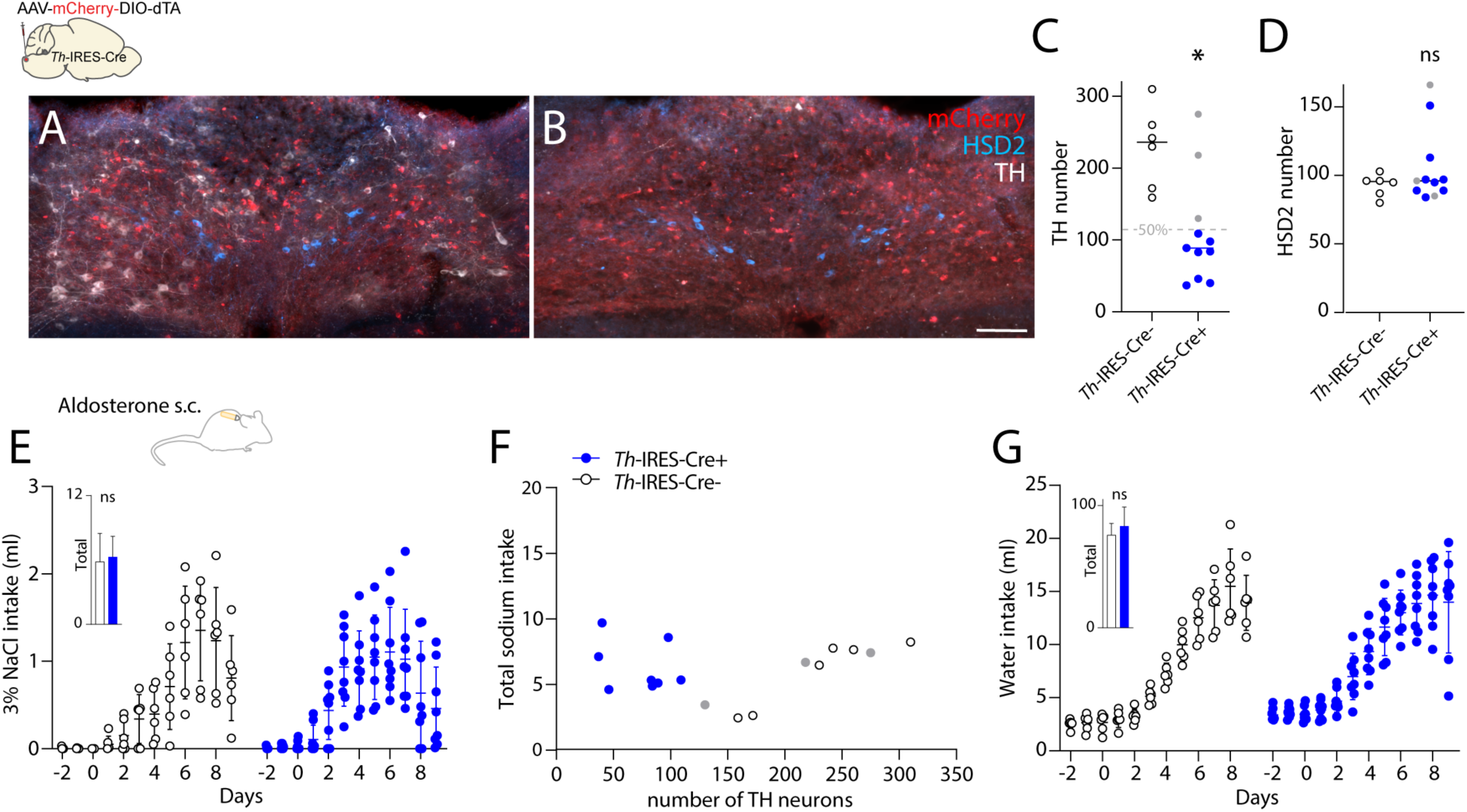
Effect of ablating NTS catecholaminergic neurons on aldosterone-induced salt and water intake. (A) After bilateral microinjections of the ablation vector (AAV-mCherry-DIO-dtA), control mice (Cre-negative littermates) had a normal distribution of TH-immunoreactive neurons (white). These and many other non-Cre expressing neurons in the injected region express mCherry. (B) Experimental mice (*Th*-IRES-Cre+) injected with the same ablation vector had few remaining TH+ neurons. In both (A) and (B), unaffected aldosterone-sensitive neurons are evident by their immunoreactivity for HSD2 (blue). (C) Genetically targeted ablation reduced the number of NTS catecholaminergic neurons (*p=0.04 relative to Cre-negative littermates, by two-tailed t-test), without altering the number of HSD2 neurons (p=0.2886). (E-G) Effects of NTS catecholaminergic neuron ablation on 3% NaCl and water intake in mice receiving a maximally effective rate of s.c. aldosterone infusion (1000 ng/h). For inset panels (E) and (G) “ns” = “not significant” (p=0.7120 and p=0.3315, respectively, relative to Cre-negative littermates).

### Water intake during peripheral infusion of aldosterone

The increase in water intake during peripheral (s.c.) infusion of aldosterone was unexpected, and to our knowledge, never previously reported. The lack of an effect with i4V aldosterone infusion and the persistence of this effect in mice lacking HSD2 neurons suggested a peripheral rather than central mechanism. Several systemic factors can increase thirst, and it is important to first differentiate “primary” polydipsia, which is thought to have a neurologic or psychiatric cause, from thirst that occurs simply due to excessive urinary loss. We began by comparing daily water intake and urine output between aldosterone-infused experimental and vehicle-infused control mice, with and without mildly restricted access to water. These mice had *ad libitum* access to normal chow and no access to saline. With unrestricted access to water, aldosterone-infused mice again drank more water (p<0.0001 relative to vehicle-infused controls), with intakes gradually rising to surpass 10 mL per day. These aldosterone-infused mice also excreted proportionally more urine (p=0.0055; **Supplemental Figure 2D–F**), with no differences in body weight change (aldosterone: −1.2 ± 0.6 g vs. vehicle: −0.2 ± 0.7 g, p=0.0769) or total food intake (aldosterone: 39.1. ± 2.6 g. vs. vehicle: 36.3 ± 2.0 g, p=0.1459). With mildly restricted access to water (5 mL per day), however, the daily urine volumes of aldosterone-infused mice remained within the range of vehicle-infused control mice (aldosterone vs. vehicle d1-7, p>0.9999). These water-restricted, aldosterone-infused mice did not lose weight (p= 0.3268), and their daily urine output did not increase until we liberalized their access to water (aldosterone d1-7 vs. d8-10; p=0.0071; aldosterone d8-10 vs vehicle d8-10, p=0.0010; **Supplemental Figures 2G–I**). These findings indicate that thirst caused by s.c. aldosterone infusion is not secondary to polyuria.

We next considered the possibility that aldosterone-infused mice were thirsty due to hypernatremia, caused by a combination of aldosterone-induced sodium intake and retention. However, the plasma sodium concentrations of aldosterone- and vehicle-infused mice were not different (p=0.9523; **Supplemental Figure 2K**). Diabetes insipidus is a well-known cause of polyuria and (secondary) polydipsia, and in a separate group of mice, we measured plasma copeptin, which is produced in direct proportion to vasopressin (the antidiuretic hormone), but there was no difference between vehicle- and aldosterone-infused mice (p=0.0907; **Supplemental Figure 2L**). Another well-known cause of polyuria and (secondary) polydipsia is hyperglycemia, and aldosteronism is associated with elevated risk of diabetes mellitus (6, 8), but blood glucose levels were not elevated in aldosterone-infused mice (p=0.1571; **Supplemental Figure 2M**). Thus, polydipsia evoked by peripheral aldosterone infusion does not result from hypernatremia, hyperglycemia, diabetes insipidus, or polyuria, and likely involves a peripheral mechanism unrelated to HSD2 neurons.

## Discussion

This study improves our understanding of sodium appetite by clarifying the dose-response relationship between aldosterone and salt intake and by identifying specific neurons required for this behavioral effect. We showed that dietary sodium deprivation elevates aldosterone, which crosses the blood-brain barrier to reach tissue concentrations within its functional range for activating the mineralocorticoid receptor. We confirmed that aldosterone activates HSD2 neurons and that activating HSD2 neurons increases sodium appetite, and we discovered that eliminating HSD2 neurons prevents aldosterone-induced salt intake. Importantly, we identified homologous HSD2 neurons in humans and pigs, suggesting that these aldosterone-sensitive, sodium-appetite-promoting neurons are an evolutionarily conserved component of the mammalian brain. Collectively, our findings highlight the HSD2 neurons as a promising target for therapies aimed at modifying human sodium intake.

### Aldosterone increases salt intake

Our findings unequivocally demonstrate that aldosterone increases sodium appetite. We found specificity (non-infused and vehicle-infused mice did not consume salt) and a clear dose-response relationship (maximal behavioral effects occurred at i4V infusion rates that precisely reproduced endogenous CSF aldosterone levels on a low-sodium diet), as well as neuroanatomical localization (i4V infusion was most potent and infusing it into the forebrain ventricular system was ineffective). This behavioral role, mediated exclusively through HSD2 neurons, complements aldosterone’s sodium-retaining effects in peripheral epithelia, making this adrenal steroid hormone a key regulator of sodium homeostasis and extracellular fluid volume.

Ours is the first study focusing specifically on this topic in mice, and the most extensive investigation of aldosterone-induced behavior in any species. The above findings update several previous misconceptions, including our own, about the possible relevance of mineralocorticoid-induced behavior in health and disease states. Knowing that aldosterone poorly penetrates the blood-brain-barrier (63, 66, 70) raised doubts as to whether it can reach brain tissue concentrations that are adequate to influence behavior (47). A different mineralocorticoid receptor agonist, deoxycorticosterone, did increase salt intake (71, 72), but this required supraphysiologic doses in rats, and subsequent investigators could not replicate this behavioral effect in mice (38, 73). Although no chronic, dose-ranging studies had been attempted with aldosterone, such findings cast doubt on the likelihood that it has a significant behavioral role.

In contrast, our central and peripheral dose-ranging tests revealed that a small but significant portion of circulating aldosterone reaches the brain. Merely a week of low-sodium diet triggered enough aldosterone production to elevate brain (CSF) concentrations into the low-nanomolar range, where it potently binds and activates the mineralocorticoid receptor. Importantly, peak salt intake occurred at i4V infusion rates that produce precisely the same range of CSF concentrations we measured in salt-hungry mice after a week of dietary sodium deprivation. Together, these findings indicate that the large, gradual rise in circulating aldosterone caused by dietary sodium deprivation is responsible for the resulting increase in sodium appetite, consistent with a previous finding in rats that i4V injection of a mineralocorticoid receptor antagonist inhibits saline intake in sodium-deprived rats (11).

Dietary sodium deprivation, along with potassium supplementation, is the most ethologically relevant paradigm for triggering aldosterone production (1, 34, 74) and sodium appetite (2, 32, 34, 51, 52, 75). These two dietary manipulations recreate the existential dilemma faced by all land-dwelling herbivores and omnivores: most foods have zero sodium and high potassium. A high-potassium diet is beneficial, but a zero-sodium diet is harmful, and eventually lethal (76), and excreting potassium on an extremely low-sodium diet requires increased aldosterone (55, 77). Increased aldosterone is an expected and life-critical response to chronic hypovolemia and gradually rising extracellular potassium level that results from chronically consuming a diet with inadequate sodium (1, 34, 78, 79). Our results confirm that, like humans and rats, mice generate very high aldosterone levels (100–1000 ng/dL) within a week after simply altering dietary sodium and potassium.

Prior to this study, it was unclear why the body would expend so much energy to hypertrophy the adrenal glomerulosa and generate such high levels of circulating aldosterone (10–20 nM), well beyond what is needed for maximal binding and activation of mineralocorticoid receptors in peripheral tissues. Based on the results of this study, we propose that the benefit of generating such high aldosterone levels, in a chronically sodium-deplete state, is that these circulating levels overcome the steep blood-brain concentration gradient (63) and sufficiently elevate brain-tissue concentrations of aldosterone to a level capable of activating the mineralocorticoid receptor in HSD2 neurons. Circulating levels at or below 50–100 ng/dL (1–3 nM) are more than sufficient to retain sodium by maximally activating peripheral mineralocorticoid receptors in HSD2-expressing epithelial cells. In contrast, activating the behavioral drive to seek and consume sodium, which is the only way to repair a chronic volume deficit, requires high levels (well above 100 ng/dL) that elevate brain-tissue concentrations and activate mineralocorticoid receptors in HSD2-expressing neurons. That is, mild elevations are adequate for sodium retention, but in a chronic zero-sodium environment, larger amounts of aldosterone are needed to increase circulating levels by roughly 10x and achieve a brain concentration adequate for activating a behavioral mode that represents the only hope for survival: seeking and consuming salt.

Like mice and rats, healthy human subjects generate circulating aldosterone levels upwards of 300 mg/dL if they are fed a low-sodium, high-potassium diet (55). Even higher aldosterone levels – more than three times this high – have been found in patients with aldosterone-secreting adenomas or chronic kidney disease (61, 62). However, besides tribes outside contemporary civilization (78) and individuals who deliberately abstain from sodium (80), most people alive today consume more than enough sodium and probably will never generate such high aldosterone levels. Thus, while it will be interesting to determine the extent to which aldosterone induces sodium appetite in sodium-deprived healthy subjects, it will be more important to investigate its behavioral effect in patients with aldosteronism.

### Cell-type-specificity of aldosterone-induced salt intake

Prior to this study, several investigators attempted to localize the site of action for mineralocorticoid-induced sodium appetite in the rat brain. In the 1960s, Wolf and colleagues focused on the hypothalamus, where large electrolytic lesions reduced several ingestive behaviors, including saline intake after peripheral injections of deoxycorticosterone (81). In the 1980s, Epstein and colleagues proposed that sodium appetite arises from a synergy between aldosterone and angiotensin II (10, 82). This synergy was thought to occur within the lamina terminalis, which lies along the anterior wall of the third ventricle and includes the subfornical organ (50). Neurons here mediate angiotensin II-induced fluid intake (46, 49, 83–87), but none express HSD2 and the mineralocorticoid receptor (29, 30). In our study, infusing aldosterone immediately upstream of this circumventricular region did not alter salt intake, replicating a previous finding in rats (11), and an additional study in rats reported that injecting angiotensin II into the lateral ventricle did not increase, and in fact reduced saline consumption when coupled with aldosterone infusion (88). In another forebrain region, the central nucleus of the amygdala, mineralocorticosteroid-induced saline intake was reduced by injecting either excitotoxins or antisense oligonucleotides designed to suppress production of the mineralocorticoid receptor (39, 41, 44, 89). Aldosterone is unlikely to act directly within this region because, as in the lamina terminalis, no cells here express HSD2 and the mineralocorticoid receptor (29, 30), but the central nucleus of the amygdala does contain neurons with direct output projections to the aldosterone-sensitive HSD2 neurons in the NTS (90).

Just as angiotensin II potently increases water intake (91), aldosterone is the most selective and individually potent stimulus for salt intake. Infusing aldosterone suppresses renin and angiotensin II (92), and the present results confirm that aldosterone is capable of inducing salt intake on its own. This singular ability of aldosterone, coupled with co-expression of the mineralocorticoid receptor and HSD2 enzyme in a single population of neurons, provided the first circumstantial evidence for involvement of these neurons in aldosterone-induced salt intake. Further circumstantial support derived from the finding that HSD2 neurons fire faster after chronic infusion of aldosterone (35), express the activity marker Fos during aldosterone infusion (shown here), and trigger saline intake when chemogenetically stimulated (35, 36). Despite the compelling circumstantial evidence, however, directly testing this hypothesis required first determining a working and physiologically relevant aldosterone dose in mice and then using cell-type-specific techniques to selectively remove these neurons and rigorously assess their role. This combined approach unambiguously showed that HSD2 neurons are necessary for aldosterone-induced sodium appetite. The HSD2 neurons are the first and, thus far, only neurons with this property.

Sodium appetite may be the primary or only central function of aldosterone and of the aldosterone-sensitive HSD2 neurons, as neither i4V infusion in rats nor chemogenetic activation of HSD2 neurons in mice altered blood pressure (12, 35). Thus, a singular hormone, aldosterone, induces a highly specific behavioral program, seeking and consuming sodium, and this behavioral switch depends upon a tiny population of neurons, numbering barely more than 100 (30). These 100 neurons represent a minority of the NTS and a tiny fraction of neurons in the brain overall – approximately 0.0001% of ∼100 million neurons in the mouse brain and 0.000001% of ∼100 billion neurons in the human brain. We are not aware of any other example of a mammalian behavior depending so completely on the integrity of so few neurons. An obvious comparator is the *Agtr1a*-expressing population of thirst-promoting neurons in the lamina terminalis, which spans the subfornical organ, median preoptic nucleus, and organum vasculosum of the lamina terminalis (87), but their total number remains unknown, and whether angiotensin-induced fluid intake entirely depends on these neurons is unclear. Another relevant comparator is the *Agrp* population of hunger-promoting neurons in the arcuate nucleus of the hypothalamus (93, 94), but these neurons are 40-fold more numerous, and both baseline food intake and (partially) fasting-induced food intake persist without them (95). Thus, the total dependency of aldosterone-induced salt intake on the integrity of HSD2 neurons may represent the most delicately cell-type-specific behavioral dependency in the mammalian brain.

Of note, while our results demonstrate an absolute requirement of HSD2 neurons for *aldosterone-induced* sodium appetite, other neurons may help increase sodium appetite in other experimental paradigms. Some neurons in the lamina terminalis may exert a natriorexigenic effect in a subacute preparation where furosemide diuresis is followed by dietary sodium deprivation for just one day (48). In this paradigm, inhibiting and ablating the HSD2 neurons had variable and subtotal effects (35, 36), and it will be important to determine the relative contributions of aldosterone and HSD2 neurons to this and other paradigms that provoke thirst and sodium appetite.

### Clinical relevance

Our finding that a singular hormone (aldosterone) activates a specific behavior (salt intake) via a distinctive cell population (HSD2 neurons in the NTS) has significant implications for patients in whom dietary sodium is already known to modify symptom severity or end-organ damage. The presence of HSD2 neurons in the human brainstem suggests translational potential, and the existence of neurons whose core function is promoting sodium appetite suggests an opportunity for targeted interventions that modify dietary sodium consumption and its associated risks and benefits in vulnerable patient populations.

Specifically, the HSD2 neurons may be a useful lever for nudging dietary sodium intake up or down, depending on the need in a specific patient. For example, salt supplementation is helpful in patients with a variety of autonomic disorders, including orthostatic hypotension, vasovagal syncope, postural orthostatic tachycardia syndrome, and neurodegenerative synucleinopathies like pure autonomic failure, Parkinson’s disease, dementia with Lewy bodies, and multiple system atrophy (80, 96, 97). Many of these patients have a deficit in retaining urinary sodium, so supplemental sodium is used to expand plasma volume. Dietary sodium is supplemented with large salt tablets, sometimes in combination with a combined mineralocorticoid-glucocorticoid receptor agonist, fludrocortisone. However, treatment benefits are frequently limited by gastrointestinal distress caused by salt tablets, by steroid side-effects, and by fludrocortisone contraindication in patients with heart failure or with supine hypertension, which is common in autonomic synucleinopathies (97). In such patients, graded stimulation of HSD2 neurons should boost an intrinsic drive to increase salt consumption gradually, in food, rather than acutely, in tablets delivering a 500–1,000 mg salt bolus. The effects of acute and chronic chemogenetic stimulation shown here in mice confirm the basic feasibility of this approach, but additional work is required to establish feasibility then evaluate efficacy and possible risks in patients.

Separately, ablating or inhibiting HSD2 neurons may help patients with salt-sensitive hypertension (98), particularly those with aldosteronism, which is a common cause of treatment-refractory hypertension (18–23). Patients with aldosteronism suffer higher rates of stroke, heart failure, and other cardiovascular complications, and their increased sodium intake explains a significant portion of this elevated risk (18–21). Cutting salt intake improved both blood pressure and mood in a 12-week trial (23), but we lack efficacious behavioral therapies for sustaining this dietary change. The cell-type-specific ablation results shown above establish the basic feasibility of eliminating HSD2 neurons in mice with excess aldosterone, but additional work is required to establish feasibility then evaluate efficacy and possible risks in patients.

Overall, an important implication of our results is that dietary sodium is a regulated variable, much like water and food. The behavioral motivation to seek and consume salt, much like the behavioral motivation to seek and consume food or water, is subject to homeostatic feedback and governed by neurons in the brain. Patients and the general public stand to benefit if we incorporate this information into clinical guidelines, nutritional recommendations, and public health policies aimed at improving the choices people make about what they eat and drink.

### New and unanswered questions and limitations

Our results include several unexpected findings that represent new opportunities for investigation. We will briefly discuss these findings and several limitations inherent to our approach.

First, despite identifying homologous neurons in the human brain, our reliance on an animal model for pharmacologic and behavioral experiments necessitates cautious extrapolation of these findings to human behavior. Further investigation in humans is warranted.

Second, peripheral infusion of aldosterone induced less salt intake than i4V infusion. This remained true even with peripheral (s.c.) infusion rates that generated CSF levels within or above the range of maximally effective i4V infusions, so this lesser sodium intake cannot be explained by inadequate aldosterone in the brain. Peripheral infusion must have had a mixed effect, stimulating sodium appetite in the brain, while at the same time inhibiting sodium appetite as a consequence of peripheral hyperaldosteronism, which was not present in mice with i4V infusion. In either paradigm, consuming salt should briefly increase plasma sodium and blood volume, which would transiently reduce HSD2 neuron activity and sodium appetite (27, 33, 34, 99–101). Aldosterone-infused mice were not hypernatremic, but peripherally infused mice may have experienced some blood volume expansion due to the combination of (central) aldosterone-induced saline intake and (peripheral) aldosterone-induced sodium retention. In contrast, i4V-infused mice likely excreted most or all of the excess saline they consumed. Blood volume expansion activates low-pressure barosensors, triggers cardiac release of natriuretic peptides, and suppresses renin and angiotensin II production. These hypothesized mechanisms are not mutually exclusive, and it will be helpful to clarify their combined impact on the activity of HSD2 neurons and other, connected neurons in the overall network controlling salt-ingestive behavior.

Third, the mechanism of increased water intake caused by peripheral aldosterone infusion remains unclear. We did not expect a thirst response because, to our knowledge, no such effect has been reported in any species. Peripheral-aldosterone-induced thirst does not appear to be mediated by well-known thirst triggers like hypernatremia, hyperglycemia, diabetes insipidus, polyuria, or hypovolemia. Similar to salt intake, daily water intake increased gradually and plateaued over the course of a week. However, stimulating HSD2 neurons had no effect on water intake and eliminating HSD2 neurons had no effect on thirst induced by peripheral infusion of aldosterone. This unexpected effect probably did not involve direct action in the brain because infusing aldosterone into the cerebral ventricles had no significant impact on water intake, and thirst appeared to increase at peripheral infusion rates below the threshold for elevating CSF levels and inducing salt intake. It would be interesting to uncover the mechanism of this unexpected effect and to determine whether it occurs in non-murine species.

Fourth, the time course of salt intake induced by chemogenetically activating HSD2 neurons was rapid, peaking the first day of infusion. In contrast, aldosterone infusion induced a slow, gradual rise in salt intake that required a week to plateau. This resembles the slow, gradual rise in saline intake following consecutive daily injections of deoxycorticosterone or i4V infusion of aldosterone in rats (11, 12, 33, 71). This slow, gradual time course, along with the lack of an acute effect of aldosterone on HSD2 neuron membrane properties in *ex vivo* tissue slices (35), appears more consistent with gradual changes in gene transcription evoked by aldosterone-bound mineralocorticoid receptors than with previous suggestions of a rapid, non-genomic effect (39, 102). Further supporting involvement of conventional mineralocorticoid receptor activity, we found maximal i4V aldosterone-induced salt intake in mice with CSF aldosterone concentrations of approximately 1 nM, the concentration at which its mineralocorticoid receptor binding and transcriptional activation is maximal (64–67). Confirming whether expression of this transcription factor in HSD2 neurons is necessary for aldosterone’s effects, and which of its transcriptomic effects are required to activate HSD2 neurons, is the subject of on-going investigation.

Fifth, it is unclear what, if any, role angiotensin II has in aldosterone-induced sodium appetite. Implanting a glass bulb to deliver angiotensin II into the dorsal hindbrain increased saline intake in rats (103). Also, the angiotensin receptor blocker losartan (20 mg/kg) reduced licking for saline after chemogenetic stimulation of HSD2 neurons in mice (35). However, eliminating angiotensin production in the liver and brain did not reduce sodium appetite in mice (54). Aldosterone infusion potently suppresses renin release and angiotensin II production (92), so the renin-angiotensin system probably plays little to no role in aldosterone-induced sodium appetite. For these reasons, we did not test the effects of infusing angiotensin or blocking renin, angiotensin converting enzyme, or angiotensin receptors. In future work, it could be interesting to explore combined effects of aldosterone and angiotensin II on HSD2 neurons *in vivo*, as well as any combined effects of aldosterone-induced activation of HDS2 neurons and angiotensin-induced activation of neurons in the lamina terminalis.

And sixth, more investigation is needed to fully understand the larger neural network that regulates sodium appetite. We proposed that HSD2 neurons integrate both synaptic and endocrine inputs, but the nature of their synaptic input and how exactly aldosterone and other endocrine signals modulate it remains unclear (35, 47, 104, 105). Also, HSD2 neurons send output to two primary targets in the brain (30, 35, 68, 69, 106). One of these targets, the fusiform subnucleus of the bed nucleus of the stria terminalis, may embody a network interaction between aldosterone-sensitive HSD2 neurons in the NTS and angiotensin-sensitive neurons in the lamina terminalis (48). The other target, the pre-locus coeruleus, may integrate input from the HSD2 neurons with input from the medial prefrontal cortex and appetite-regulating neurons in the arcuate and paraventricular hypothalamic nuclei (106). Fully understanding this aldosterone-modulated network will require detailed investigation into the connections and functions of each connected node.

## Conclusion

The HSD2 neurons are necessary for aldosterone-induced sodium appetite. Aldosterone activates these neurons, activating them increases salt intake, and eliminating them prevents aldosterone-induced salt intake. Identifying the specific cells necessary for aldosterone-induced salt intake improves our understanding of the neural circuitry controlling sodium appetite and offers a promising target for therapeutic strategies aimed at boosting sodium appetite in hypovolemic patients and mitigating excessive salt intake to improve cardiovascular health of patients with aldosteronism.

## Role of Authors

MH supplied human and porcine tissue, and LP performed immunohistology, *in situ* hybridization, and slide-scanning microscopy, as well as morphological analysis of human HSD2 neurons and early draft illustrations. SG performed all mouse experiments. SG collected, organized, and analyzed data. MCM performed chemogenetic CNO infusion experiments with SG and with supervision from JMR. CJAM counted cells in mouse tissue. JCG supervised the project, planned experiments with SG, and analyzed all results with SG. SG and JCG drafted and edited the manuscript and figures together. All authors read, commented on, and approved the final manuscript.

## Acknowledgements

This work was supported by generous funds from the University of Iowa Aging, Mind, and Brian Initiative (AMBI), laboratory startup funds from the University of Iowa Department of Neurology, a Williams-Cannon Fellowship from the Iowa Neuroscience Institute, and an Early-Stage Investigator Award from the Carver Trust. We thank Andrew Thatcher for assistance with histology and cell-counting and Richard Palmiter for generating and sharing *Hsd11b2*-Cre mice. We also thank the University of Iowa Office of Animal Resources and Iowa Neuroscience Institute for providing room space for continuous BioDAQ recordings in our mouse barrier facility, and the University of Iowa Metabolic Phenotyping Core for facilitating experiments in metabolic cages. Finally, we thank Jady Tolda for proofreading this manuscript.

## Conflict of interest statement

The authors have declared that no conflict of interest exists.

## Funding support

Laboratory startup funds (University of Iowa Department of Neurology) University of Iowa Aging, Mind, and Brian Initiative Williams-Cannon Fellowship (Iowa Neuroscience Institute) Early-Stage Investigator Award (Carver Trust & Iowa Neuroscience Institute)

## Notes

### Competing Interest Statement

The authors have declared no competing interest.

