## Supplemental Figures 1 & 2 for "HSD2 neurons are evolutionarily conserved and required for aldosterone-induced salt appetite"

### Supplemental Figures and Legends

**Supplemental Figure 1.** HSD2 neurons in several mammalian species have a similar location, distribution, and appearance. (A) HSD2-immunoreactive neurons in rat NTS (brown DAB immunohistochemical labeling for HSD2 in a horizontal tissue section, with thionin counterstain), as described in (29). (B) HSD2-immunoreactive neurons in a mouse from this study (inverted grayscale image of Cy5 labeling; coronal tissue section through the caudal medulla of an aldosterone-infused mouse). (C) HSD2-immunoreactive neurons in pig NTS (NiDAB-immunohistochemical labeling in a coronal tissue section through the medulla). (D) HSD2-immunoreactive neurons in human NTS (inverted grayscale image of Cy3 labeling in a transaxial tissue section through the caudal medulla of case MH010, also shown in Figure 1E). Scale bar in each panel is 100 µm.

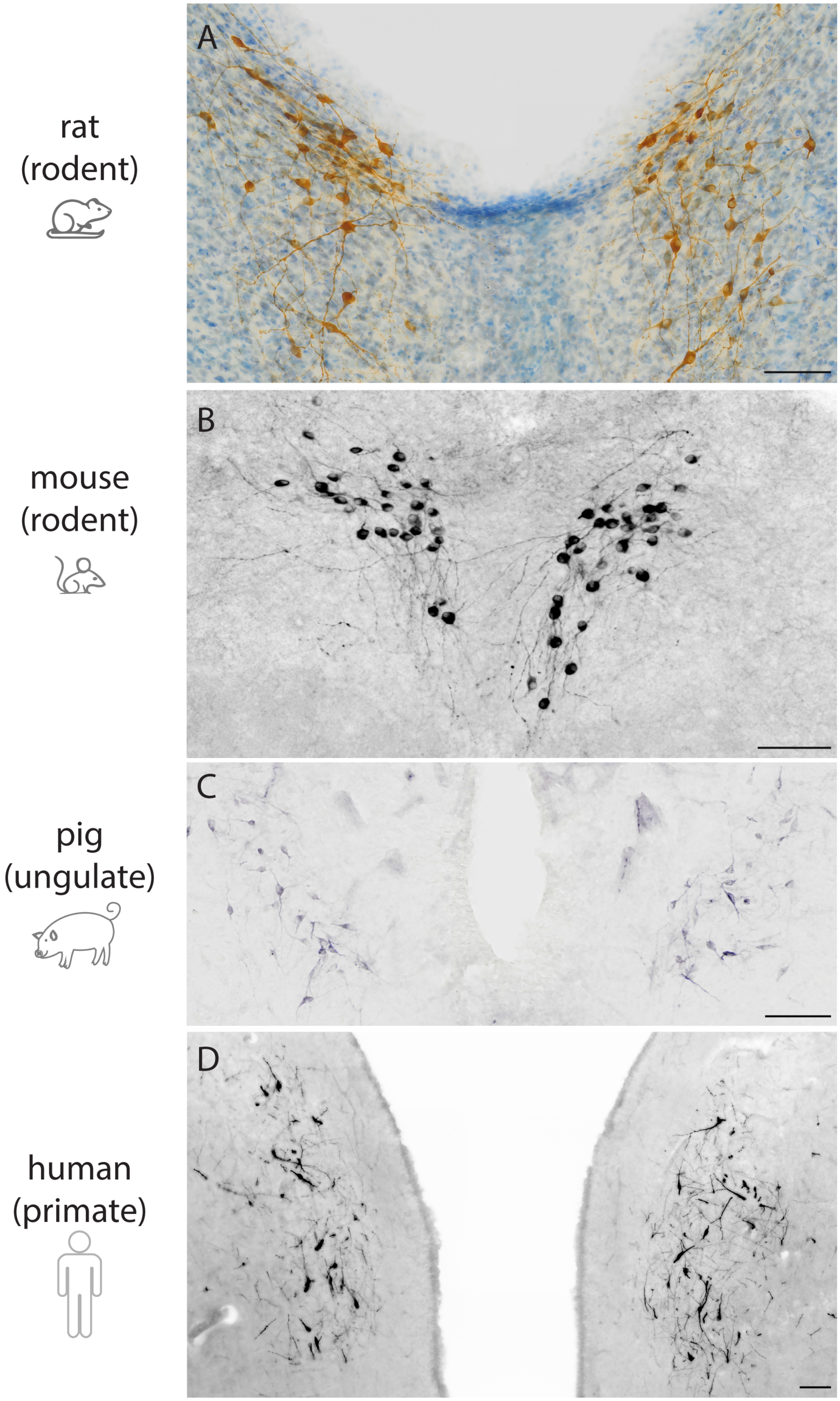

rat
(rodent)
mouse
(rodent)
pig
(ungulate)
human
(primate)
A
B
C
D

**Supplemental Figure 2.** Water intake, urine output, and blood measurements during s.c. infusion of aldosterone (1000 ng/h) or vehicle. (A-B) Daily and total water intake in mice treated with s.c. aldosterone or vehicle and given *ad libitum* access to water. Aldosterone increased total water intake ($p<0.0001$). (C-D) Daily and total urinary volume in mice treated with s.c. aldosterone or vehicle and given *ad libitum* access to water. Urine output was higher in animals treated with aldosterone ($p=0.0055$). (E) Scatterplot compares daily water intake and urine output volumes across all mice treated with s.c. aldosterone or vehicle and given *ad libitum* access to water. (F-G) Daily and total water intake in mice treated with s.c. aldosterone or vehicle and during and after mild water restriction (5 mL/d for 7d). Water intake increased in aldosterone-infused mice only when provided unrestricted access to water on days 8–10 (aldosterone vs. vehicle; $p = 0.0014$; aldosterone d1-7 vs. aldosterone 1-7d. vs. aldosterone 1-8d; $p = 0.0025$). (H-I) Daily and total urinary volume in mice treated with s.c. aldosterone or vehicle during and after mild water restriction (5 mL/d for 7d). There was no increase in urine output in the period that water is restricted (blue shade; $p > 0.9999$). When water intake increased in aldosterone-infused mice (d8–10), urine output increased as well (aldosterone 1-7d vs. 8-10d; $p =0.0025$; aldosterone 8-10d vs vehicle 8-10d, $p = 0.0014$). (J) Scatterplot compares daily water intake and urine output volumes across all mice treated with s.c. aldosterone or vehicle and during and after mild water restriction (5 mL/d for 7d). (K) Plasma sodium (mEq/L), (L) plasma copeptin (ng/dL), and (M) blood glucose (mg/dL) were not different between s.c. aldosterone- and vehicle-infused mice ($p=0.9523$; $p=0.0907$; $p=0.1571$, respectively).

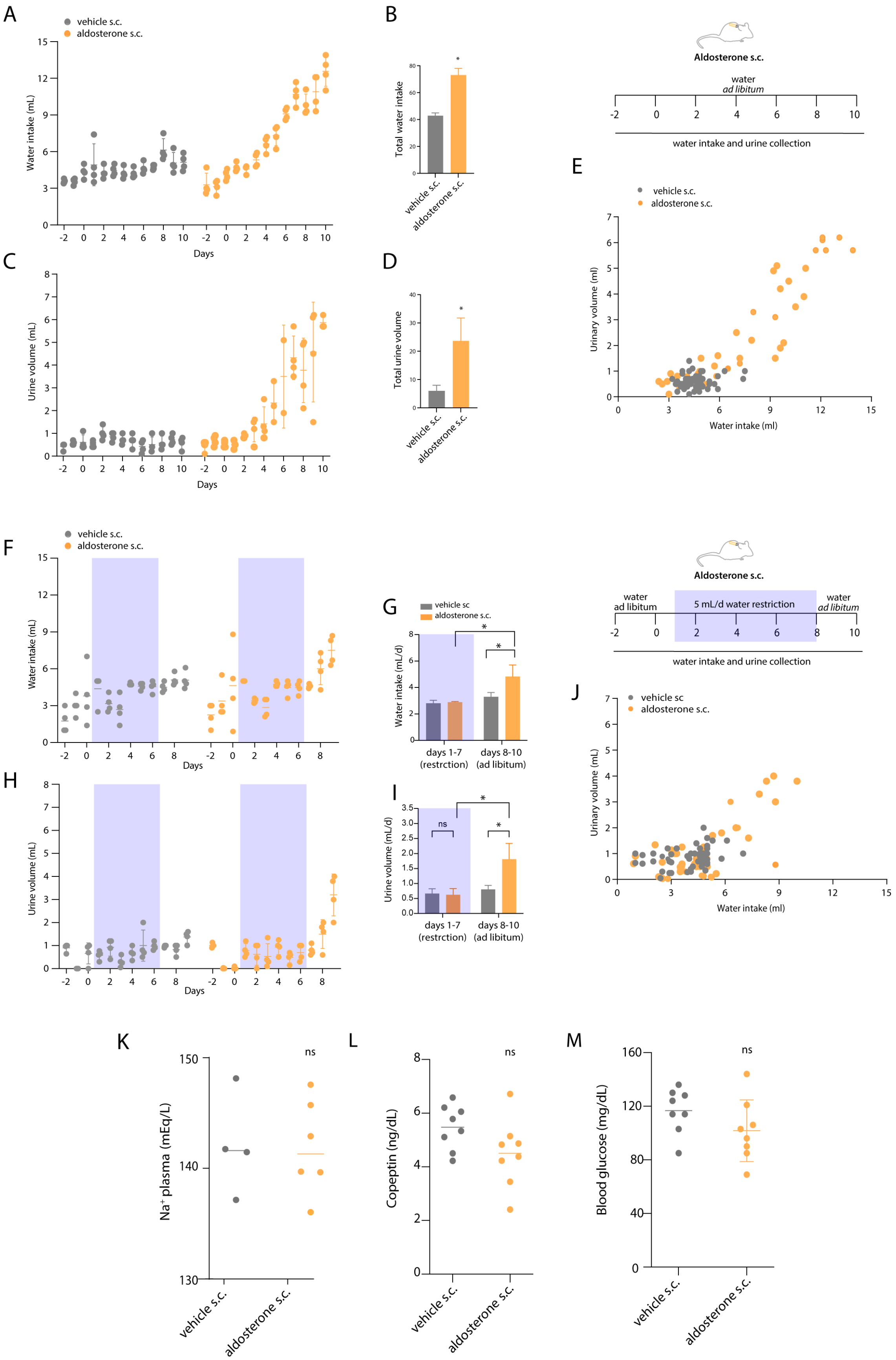
A
vehicle s.c.
aldosterone s.c.
Water intake (mL)
Days
B
Total water intake
vehicle s.c.
aldosterone s.c.
Aldosterone s.c.
water
ad libitum
water intake and urine collection
C
Urine volume (mL)
Days
D
Total urine volume
E
Urinary volume (ml)
Water intake (ml)
F
Water intake (mL)
Days
G
vehicle sc
aldosterone s.c.
Water intake (mL/d)
days 1-7 (restrction)
days 8-10 (ad libitum)
5 mL/d water restriction
water ad libitum
H
Urine volume (mL)
I
Urine volume (mL/d)
ns
J
vehicle sc
aldosterone s.c.
Urinary volume (mL)
Water intake (ml)
K
Na+ plasma (mEq/L)
L
Copeptin (ng/dL)
M
Blood glucose (mg/dL)
